# Microbial Energy Metabolism Fuels a CSF2-dependent Intestinal Macrophage Niche within Tertiary Lymphoid Organs

**DOI:** 10.1101/2022.03.23.485563

**Authors:** Pailin Chiaranunt, Kyle Burrows, Louis Ngai, Eric Y Cao, Siu Ling Tai, Helen Liang, Homaira Hamidzada, Anthony Wong, Meggie Kuypers, Tijana Despot, Abdul Momen, Sung Min Lim, Thierry Malleavey, Tyrrell Conway, Hiromi Imamura, Slava Epelman, Arthur Mortha

## Abstract

Maintaining intestinal macrophage (MP) heterogeneity is critical to ensure tissue homeostasis and host defense. The gut microbiota and host factors are thought to synergistically shape colonic MP development, although there remains a fundamental gap in our understanding of the details of such collaboration. Here, we report tertiary lymphoid organs (TLOs), enriched in group 3 innate lymphoid cells (ILC3s), as a microbiota-operated intestinal niche for the development of monocyte-derived MPs. ILC3-derived colony stimulating factor 2 (CSF2) serves as a developmental and functional determinant for MPs and required microbe-derived extracellular adenosine 5’-triphosphate (ATP) as a trigger. Microbial communities rich in extracellular ATP promoted MP turnover via ILC3 activity in an NLRP3-dependent fashion. Single cell RNA-sequencing of MPs revealed unique TLO-associated, CSF2-dependent MP populations critical for anti-microbial defense against enteric infection. Collectively, these findings describe a fundamental framework that constitutes an intestinal MP niche fueled by microbial energy metabolism.

## Introduction

Intestinal macrophages (MPs) represent a large proportion of the innate immune system in the gut and are critical mediators of host defense and tissue homeostasis. Research in the past decade has revealed the extensive heterogeneity in these cells, from their differential ontogeny to their location-specific divisions of labor (Chiaranunt et al., 2021). However, mechanisms regulating MP heterogeneity in the intestinal lamina propria (LP) remain enigmatic, particularly regarding the involvement of microanatomic environments that balance the abundance of MPs involved in host defense and MPs regulating tissue homeostasis.

Further complicating this matter are the classification strategies for intestinal MPs. As in other organs, intestinal MP subpopulations can be distinguished based on their expression of the markers Tim-4 and CCR2 to denote fetal-derived long-lived, self-renewing, tissue resident cells and monocyte-derived ones, respectively (Dick et al., 2022; Kang et al., 2020). Others have demarcated gut MPs using Tim-4 and CD4 into 3 subpopulations: long-lived Tim-4^+^CD4^+^ MPs, Tim-4^-^CD4^+^ MPs with slow monocytic turnover, and Tim-4^-^CD4^-^ MPs with rapid turnover (Liu et al., 2019; Shaw et al., 2018). Different populations of self-maintaining gut-resident MPs associate with neurons, vasculature, and other immune cells and reside in distinct regions of the gut, where they adopt transcriptional profiles and functions tailored to these microenvironments (De Schepper et al., 2018; Matheis et al., 2020; Muller et al., 2014). Unlike in most other organs, MPs within these intestinal microenvironments integrate signals derived from the commensal microflora into their homeostatic function (Mortha et al., 2014; Muller et al., 2014).

Several reports suggest that microbial metabolites affect intestinal MP function. For example, polysaccharides produced by *Helicobacter hepaticus* and commensal bacteria-derived short-chain fatty acids (SCFAs) were shown to promote tolerogenic MPs (Chang et al., 2014; Danne et al., 2017; Schulthess et al., 2019). Bacteria-metabolized dietary tryptophan controls monocyte differentiation in an aryl hydrocarbon receptor (AhR)- dependent manner (Goudot *et al*., 2017). Colonization with the protozoan commensal *Tritrichomonas musculis* (*T.mu*) was recently shown to induce monocyte infiltration in the gut by increasing luminal extracellular adenosine 5’-triphosphate (ATP) levels (Chiaranunt *et al*., 2022). This raises the question of whether a ubiquitously produced metabolite across microbial kingdoms may serve as a molecular motif to determine MP heterogeneity.

Microbiota and host-derived factors are proposed to collaborate in orchestrating gut MP composition and function. Deficiency in the host-derived myeloid growth factor colony stimulating factor 1 (CSF1) results in a systemic decrease in MPs, with a less pronounced effect on the intestinal tract, suggesting compensatory growth factors (Dai *et al*., 2002; Sehgal *et al*., 2018; Witmer-Pack *et al*., 1993). Interleukin (IL)-34, transforming growth factor *β* (TGF*β*), or colony stimulating factor 2 (CSF2), have been reported to affect MP development in many organs, including the intestinal tract (Greter *et al*., 2012; Guilliams *et al*., 2013; Schridde *et al*., 2017). In the gut, tissue-resident group 3 innate lymphoid cells (ILC3s) produce large quantities of CSF2 in a microbiota-dependent manner within intestinal tertiary lymphoid organs (TLOs), such as cryptopatches and isolated lymphoid follicles (Mortha *et al*., 2014). *Csf2*^-/-^ mice displayed a partial reduction in intestinal MPs, suggesting that MP development may in part depend on this growth factor (Mortha *et al*., 2014). CSF2 has recently been shown to license the effector profile of MPs in the inflamed brain, implicating an impact on MP function in addition to development (Amorim *et al*., 2022). Whether these observations on CSF2 extend to MP development and function in the intestine remain unknown.

Here, we report a molecular and spatial framework governing the collaboration between host and microbiota that regulates colonic MP heterogeneity. Combining fate-mapping models, immunofluorescence assays, microsurgical dissection, and single cell RNA-sequencing (scRNA-Seq) analysis, we identified TLOs as a supporting niche for the developmental and functional programming of monocyte-derived TLO-associated Tim-4^-^ CD4^-^ MPs. Using adoptive transfer experiments and mono-colonization of germ-free mice, we demonstrated that microbe-derived extracellular ATP serves as a driver of CSF2 production by ILC3s in an NLRP3-dependent fashion to induce the monocyte to MP transition. TLO-associated MPs expressed distinct genes, displayed high metabolic demand, and followed an alternative differentiation pathway compared to LP-resident MPs. Development of TLO-associated MPs was dependent on CSF2 and protected the host against enteric bacterial infections. Collectively, our findings identify a molecular and spatial framework for the location-specific differentiation of colonic MPs, centered around a microbiota-fueled, ILC3-driven and CSF2-dependent axis that integrates signals indicative of microbial energy into the heterogeneity of colonic MPs.

## Results

### Colonic monocyte-derived MPs require CSF2

Tissue-resident MPs in extra-intestinal organs group into monocyte- or fetal-derived subpopulations based on cross-organ conserved expression of the markers CCR2, Tim-4, LYVE1, and MHCII (Dick *et al*., 2022). In the gut, MPs have been classified by Tim-4 and CD4 expression and the ‘monocyte waterfall’ gating strategy (Bain et al., 2014; Shaw *et al*., 2018). To consolidate these various gating strategies, we performed an unbiased t-distributed stochastic neighbor embedding (t-SNE) dimensionality reduction on colonic lamina propria (LP) CD64^+^ CD11b^+^ cells. Distinct and partial overlap in the expression of Tim-4, CD4, CCR2, MHCII, and Ly6C aligned with the previous classification into Tim-4^+^CD4^+^, Tim-4^-^CD4^+^, and Tim-4^-^CD4^-^ MPs. Tim-4^-^CD4^-^ MPs were further delineated into Ly6C^+^, CCR2^+^, and CCR2^-^ MPs, suggesting a developmental relationship to monocytes (**Fig 1A, Fig S1A,B**)(Dick *et al*., 2022; Shaw *et al*., 2018). Accordingly, Tim-4^-^CD4^-^ MPs were significantly depleted in *Ccr2*^-/-^ mice, leaving Tim-4^+^ MPs as the majority of colonic MPs in these mice (**Fig 1B**). Previous investigations into *Csf2^-/-^* mice did not use this detailed classification of MPs, prompting us to revisit the requirements for CSF2 on gut MP heterogeneity (Mortha *et al*., 2014). Analysis of MPs in the colonic LP of *Csf2*^-/-^ mice revealed an elevated abundance of Ly6C^hi^ monocytes compared to WT or *Ccr2*^-/-^ mice, implicating a developmental blockade on the transition from monocytes to MPs (**Fig S1C**). CCR2^+^ and CCR2^-^ Tim-4^-^CD4^-^ MPs and Tim-4^-^CD4^+^ MPs were reduced in *Csf2*^-/-^ mice, further implicating their developmental relation to monocytes (**Fig 1B, Fig S1C**). Tim-4^+^ MPs were partially affected in *Csf2^-/-^* mice, suggesting that all MP subpopulations variably depend on this growth factor (**Fig 1B, Fig S1C**).

**Figure 1.**
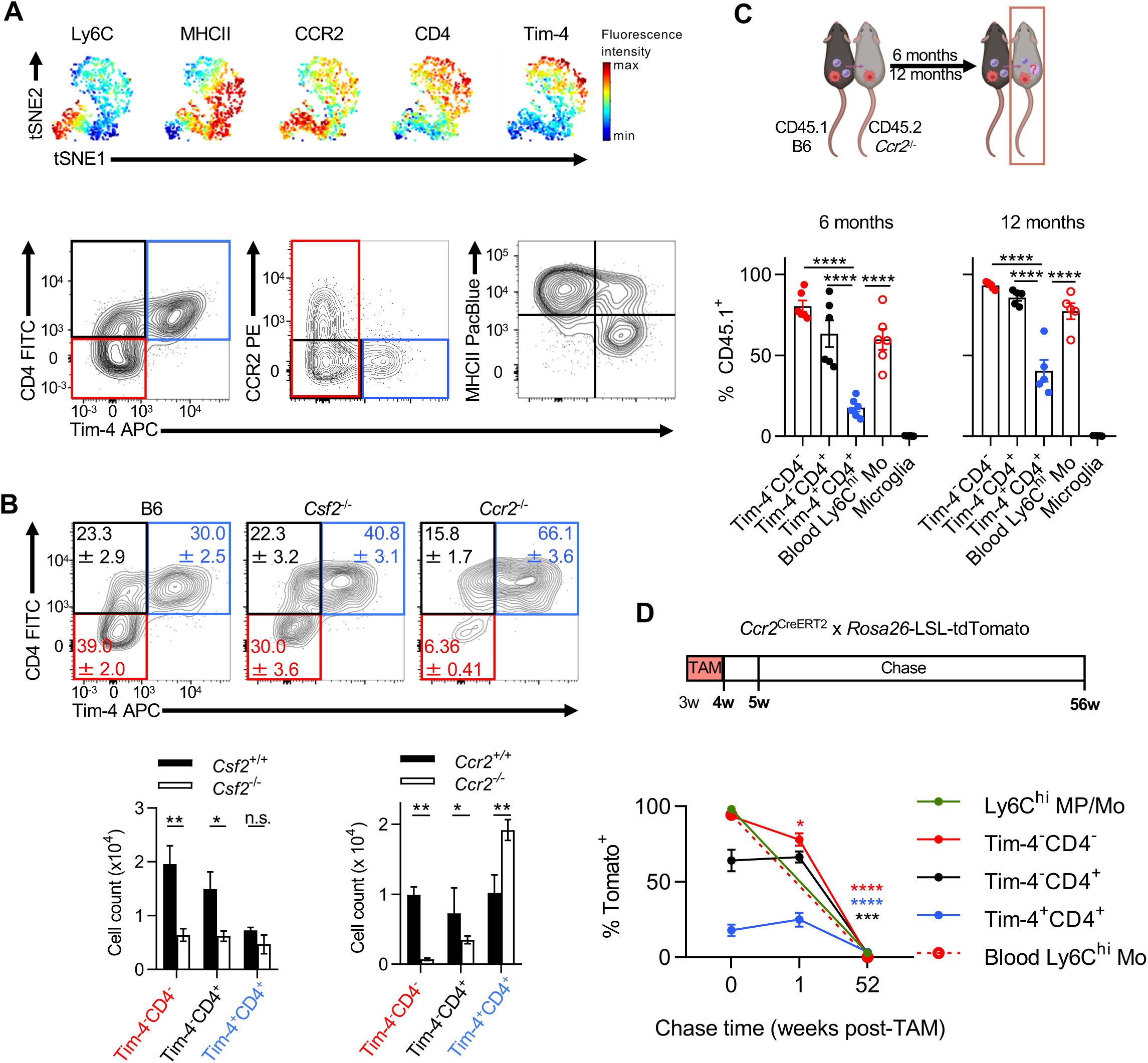
Developmental requirements and kinetics of colonic lamina propria MPs. (A) Representative flow cytometry analysis using (top) unbiased t-SNE dimensionality reduction of colonic CD64^+^ CD11b^+^ cells showing expression of common MP markers, or (bottom) Tim-4/CD4, Tim-4/CCR2, and Tim-4/MHCII classification strategies. (B) Contour plots (top) and quantification (bottom) of colonic MPs in B6, *Csf2*^-/-^, and *Ccr2*^-/-^ mice. (C) CD45.1 B6 and CD45.2 *Ccr2*^-/-^ female parabiotic pairs (top) were analyzed at 6 months (bottom left) or 1 year (bottom right) after surgery. Chimerism was quantified for colonic MP populations, blood monocytes and microglia. (D) *Ccr2*^CreERT2^ x *Rosa26*^td^ 3-week-old mice were fed tamoxifen-containing chow for 1 week, then assessed for loss of Tomato labeling in colonic MP populations at 0, 1, and 52 weeks post withdrawal of tamoxifen. Significance is calculated for each group comparing to previous timepoint. Data shown in (A) and (B) are representative of at least three independent experiments with at least three mice per group per experiment. Data shown in (C) and (D) are from two independent experiments. Two-way ANOVA (for (B and D)) or one-way ANOVA (for (C)) with post-hoc Tukey’s test was performed; *p < 0.05, **p < 0.01, ***p < 0.001, ****p < 0.0001; n.s., not significant.

Tim-4^+^ MPs are a long-lived, fetal-derived population with minimal replacement by monocytes. It has further been proposed that the expression of Tim-4 may reflect long-term residency within a tissue following differentiation (Bleriot et al., 2020; De Schepper *et al*., 2018; Scott et al., 2016; Shaw *et al*., 2018; Theurl et al., 2016). To determine the long-term MP turnover by infiltrating monocytes, we performed parabiosis of CD45.1 C57BL/6 and CD45.2 *Ccr2*^-/-^ mice to assess the chimerism of each MP subpopulation after 6 or 12 months. As expected, *Ccr2*^-/-^ parabionts showed CD45.1 frequencies comparable to blood monocytes in both the colonic Tim-4^-^CD4^-^ MP s and Tim-4^-^CD4^+^ MP compartments (**Fig 1C**). Surprisingly, Tim-4^+^ MPs were also replaced by donor CD45.1 monocytes, albeit at a slower rate, suggesting a homeostatic contribution of monocytes to the maintenance of Tim-4^+^ MPs (**Fig 1C**). To confirm our results, we employed a tamoxifen-inducible fate-mapping model. Tamoxifen-containing chow was provided to *Ccr2*^CreERT2^ x *Rosa26*-LSL-tdTomato (*Rosa26*^td^) mice during a 1-week pulse phase, followed by a chase period with normal chow for either 1 or 52 week(s). Loss of Tomato labeling was measured in each MP subpopulation to assess the replacement of MPs by newly infiltrated monocytes. Tamoxifen administration labelled ∼94% of all blood Ly6C^hi^ monocytes within the 1-week pulse phase. The induced Tomato label was absent in monocytes after 52 weeks (**Fig 1D**). Colonic Ly6C^+^ monocytes and Tim-4^-^CD4^-^ MPs showed labeling efficiency similar to blood monocytes, while Tim-4^-^CD4^+^ MPs displayed partial Tomato labeling (64%). Surprisingly, ∼17% of all Tim-4^+^ MPs were labeled after the 1-week pulse phase, suggesting a possible contribution of monocytes to this population during tamoxifen administration (**Fig 1D**). After the first 7 days of the chase period, Tim-4^-^CD4^-^ MPs (containing Ly6C^+^ and CCR2^+^ cells) displayed replacement by bone marrow-derived (BM) monocytes, as indicated by the loss of Tomato labelling (**Fig 1D**). Tim-4^-^CD4^+^ MPs and Tim-4^+^ MPs did not show signs of replacement, suggesting a slower replacement rate in line with our parabiosis experiments (**Fig 1C, 1D**). After the 52-week chase period, all MP subpopulations lost the Tomato label (**Fig 1D**). These results indicate that all colonic MPs share a monocytic origin, with Tim-4^-^CD4^-^ MPs showing the fastest replacement and strongest reliance on CSF2.

### Diversified microbiotas promote the accumulation of Tim-4^-^CD4^-^ MPs

Tissue-resident Tim-4^+^ MPs dominate the MP pool during embryogenesis and are found in all tissues at early time points of fetal development, while Tim-4^-^ MPs postnatally arise from BM monocytes (Dick *et al*., 2022). In the gut, this development requires the microbiota (Bain *et al*., 2014; Shaw *et al*., 2018). Our fate-mapping and parabiosis data show that colonic MP subpopulations display different rates of monocyte replacement, suggesting distinct appearances of the MP subpopulations in the neonatal and adult intestine. To delineate the developmental kinetics of colonic MPs, we assessed the composition of MPs starting in the neonatal colon, tracking along the first weeks of life until adulthood. In line with previous reports, embryonically-derived Tim-4^+^CD4^+^ MPs primarily comprise the colons of newborn mice, followed by a significantly increased abundance of Tim-4^-^CD4^-^ MPs at 3-4 weeks of age. This time corresponds to weaning and the establishment of a diversified microbiota (**Fig 2A**)(Knoop *et al*., 2017). By 8-12 weeks after birth, Tim-4^-^CD4^-^ MPs comprise the majority of colonic MPs, implicating a microbiota-driven adaptation of the MP pool. These adaptations in gut MPs mirror previously reported observations of intestinal CSF2 production, which similarly increased until week 8 in a microbiota-dependent fashion (Mortha *et al*., 2014). Depletion of the microbiota using broad-spectrum antibiotics in adult mice shifted the MP pool in favor of Tim-4^+^CD4^+^ MPs, confirming a requirement of the microbiota in regulating MP composition (**Fig 2B**). In contrast, reconstituting germ-free mice with an adult SPF microbiota increased Tim-4^-^CD4^-^ MPs at the expense of Tim-4^+^CD4^+^ MPs (**Fig 2C**). Colonization of adult SPF mice with the protozoan commensal *T. mu* further increased the abundance of Tim-4^-^CD4^-^ MPs (**Fig 2D**). A comparable shift towards Tim-4^-^CD4^-^ MPs was also observed when analyzing “re-wilded” mice, i.e. ex-SPF mice that were colonized with the microbiota found in pet-store mice (**Fig 2E**). These findings suggest that the increase in Tim-4^-^CD4^-^ MPs in the colon may be due to an increase in microbiota-driven monocyte replacement. To track the rate of monocyte replacement in the colon, we labeled all *Cx3cr1*-expressing cells in tamoxifen-inducible *Cx3cr1*^CreERT2^ x *Rosa26*^td^ mice and followed the loss of Tomato labeling in each MP population after colonization with *T. mu* (**Fig 2F**). Compared to uncolonized littermate controls, *T. mu*-colonized mice showed significantly reduced percentages of Tomato^+^ cells particularly in Tim-4^-^CD4^-^ MPs, suggesting an increased replacement of these cells by Tomato^-^ monocytes (**Fig 2G**). In support of our fate mapping and parabiosis data, monocyte replacement was also elevated in Tim-4^-^CD4^+^ MPs and Tim-4^+^CD4^+^ MPs. Collectively, our data demonstrate that diversifying the gut microbiota promotes MP replacement by monocytes and the accumulation of Tim-4^-^CD4^-^ MPs. Notably, colonization with *T. mu* has previously been found to increase the production of CSF2 by ILC3s in the intestinal tract, implicating a role for ILC3s in the elevated MP replacement rates (Chudnovskiy *et al*., 2016).

**Figure 2.**
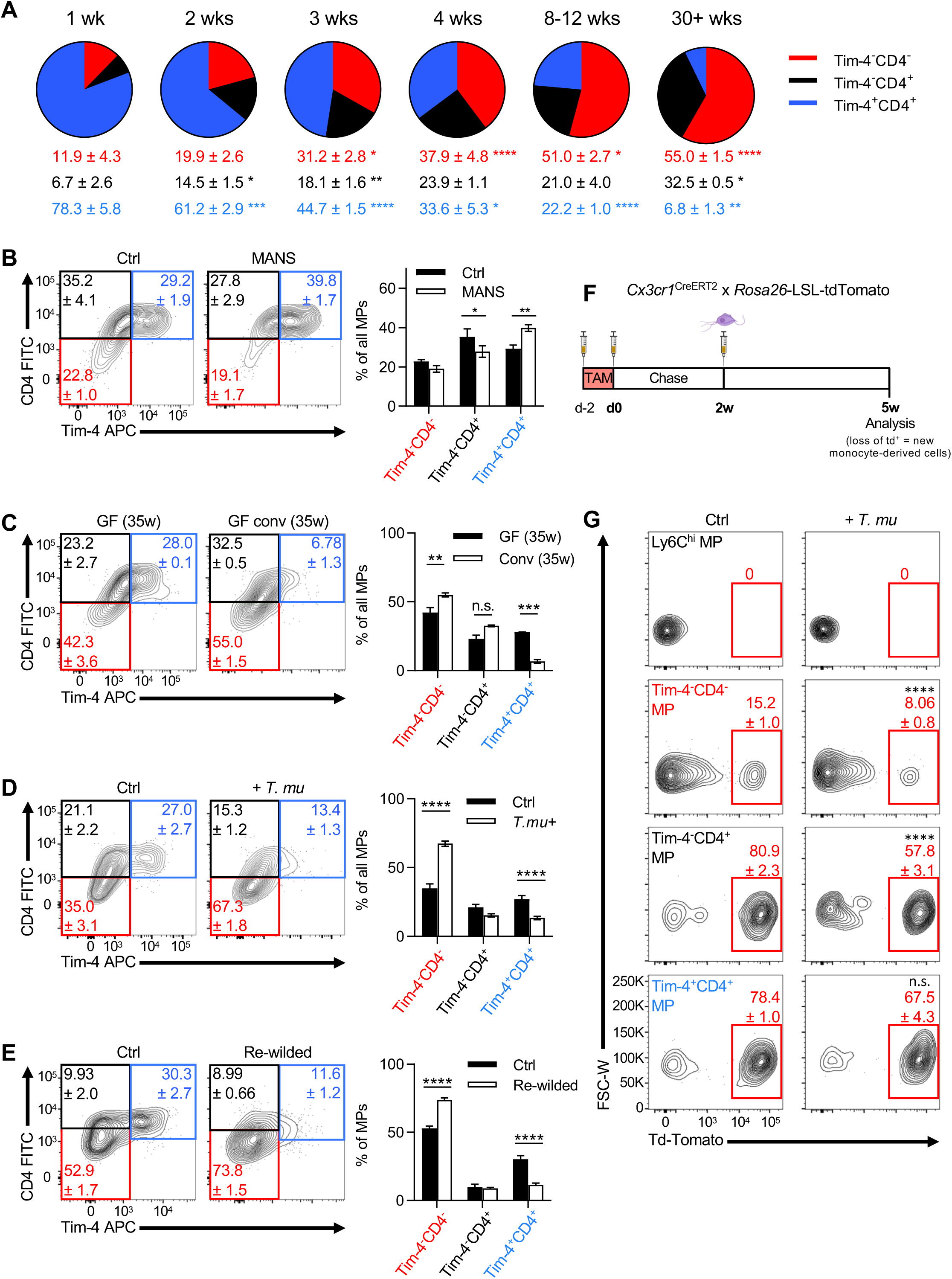
The development of colonic Tim-4^-^CD4^-^ MPs requires an intact and diversified microbiota. Colonic MP composition in (A) B6 mice across various ages, (B) B6 mice either left untreated or treated with broad-spectrum antibiotics (metronidazole, ampicillin, neomycin, streptomycin; MANS), (C) germ-free (GF) mice or GF mice conventionalized with SPF microbiota, (D) B6 mice colonized with *T. mu*, and (E) re-wilded B6 mice reconstituted with microbiota derived from pet-store mice. (F) *Ccr2*^CreERT2^ x *Rosa26*^td^ mice were injected with tamoxifen, and colonized with *T. mu* after 2 weeks via oral gavage. Colonic MPs were analyzed for Tomato label 3 weeks later. (G) Representative flow cytometry plots of mice in (F) with quantification of % Tomato^+^ cells in each colonic MP compartment. Data shown are representative of at least three independent experiments with at least three mice per group per experiment. Multiple unpaired t-tests with two-stage Benjamini, Krieger, & Yekutieli FDR test was performed for (A), *Q* = 5%, reporting q-values; each time point compared to previous time point. Two-way ANOVA with post-hoc Sidak’s multiple comparisons test was performed for (B-E), unpaired Student’s t-test was performed for each group in (G). *p < 0.05, **p < 0.01, ***p < 0.001, ****p<0.0001; n.s., not significant.

### Microbial extracellular ATP regulates MP composition and CSF2 production

Diversification of the microbiota manifests multiple new features within the microbial community, including adaptations to nutrients and synthesis of different metabolites (Blaut and Clavel, 2007). Such changes may not apply to individual microbial species but rather reflect a feature of complex community interactions (Patnode *et al*., 2021). This raises the possibility for a conserved ubiquitously produced metabolite across all living microbes that indicates microbial vitality, but at the same time, serves as a molecular motif for immune recognition and activation that can indirectly impact MP homeostasis. ATP is one such metabolite capable of promoting intestinal immunity (Atarashi et al., 2008). We recently demonstrated that colonization with the protozoan commensal *T. mu* induces immune activation in the colon, including increased monocyte infiltration, regulated by elevated intestinal extracellular ATP (ATP^ex^) levels and inflammasome activation (Chiaranunt *et al*., 2022). ATP^ex^ is a common danger associated molecular pattern (DAMP) that correlates with the presence of the microbiota and subsequently regulates local adaptive immune cells through P2X7R-dependent recognition by CD11c^+^ myeloid cells (Atarashi *et al*., 2008; Perruzza et al., 2017). Thus, we asked whether ATP^ex^ might serve as a rheostatic indicator of the microbiota, capable of regulating colonic MP composition. In line with previous studies, we first confirmed that levels of ATP^ex^ in the gut lumen corresponded to abundance of the microbiota by comparing fecal ATP^ex^ in SPF, antibiotics-treated, and germ-free mice (**Fig 3A**). ATP was shown to be released by multiple bacterial species through an unknown mechanism while undergoing cellular respiration during the growth phase *in vitro* to prolong their stationary survival (Mempin et al., 2013). To specifically investigate whether bacteria-derived ATP could regulate MPs in the gut, we utilized a mutant strain of commensal *E. coli* MG1655 that lacks the operons encoding the ATP synthase subunits A-G (*Δ*(*atpA-atpG*)) (Jones *et al*., 2007). In contrast to wild-type *E. coli* or a mutant lacking nitrate reductase genes (*ΔnarG ΔnarZ Δ*(*napD-napA*)), the ATPase-deficient mutant was unable to secrete ATP during growth *in vitro*, confirming an ATPase-dependent increase in ATP^ex^ (**Fig S2A,B**). Transformation of the Förster resonance energy transfer (FRET)-type ATP biosensor ATeam 3.10 into these *E. coli* strains enabled the quantification of intracellular ATP (ATP^int^) levels (**Fig S2C**) (Imamura et al., 2009). Accordingly, less pronounced ATP^int^ levels were observed in the ATPase-deficient *E. coli in vitro*, confirming its metabolic impairment and release of ATP (**Fig S2D,E**).

**Figure 3.**
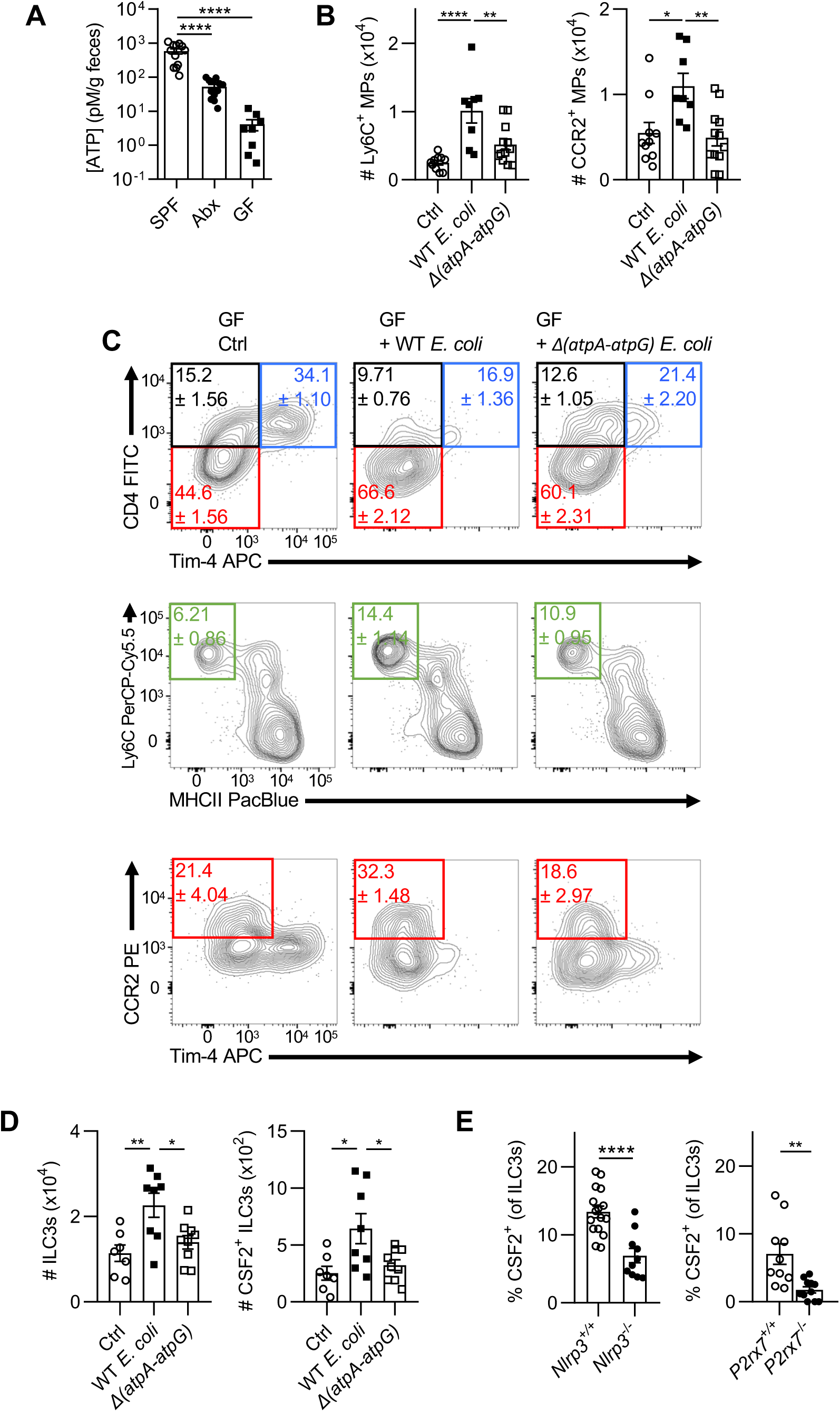
Microbial ATP regulates MP composition and drives CSF2 production by ILC3s. (A) Extracellular ATP levels in supernatants extracted from freshly isolated fecal pellets in SPF, MANS-treated SPF, and GF mice. (B-D) GF mice were orally gavaged with WT or *Δ(atpA-atpG) Strr E. coli* MG1655 and analyzed 1 week later. (B) Quantifications of Ly6C^+^ and CCR2^+^ MPs. (C) Representative flow cytometry plots show colonic monocyte and MP composition. (D) Quantifications of total ILC3s and CSF2- producing ILC3s in the colon. (E) Percentage of CSF2-producing colonic ILC3s in *Nlrp3*^-/-^ and *P2rx7*^-/-^ mice with respective littermate controls. Data shown are representative of at least three independent experiments with at least three mice per group per experiment. One-way ANOVA with post-hoc Tukey’s test (A-D) or Student’s t-test (E) was performed; *p < 0.05, **p < 0.01, ***p < 0.001, ****p<0.0001; n.s., not significant.

To determine whether an impaired ATP metabolism in commensals impacts MP heterogeneity in the gut, GF mice were mono-colonized with either control or *Δ*(*atpA-atpG*) *E. coli*. The colonic MP composition was assessed 7 days after engraftment to avoid confounding effects of adaptive immune cells on the colonizing microbes (Hapfelmeier *et al*., 2010; Macpherson and Uhr, 2004). Despite equal colonization, *Δ*(*atpA-atpG*) *E. coli* produced less ATP^int^ at the time of analysis compared to the wild-type control strain (**Fig S2F,G**). Accordingly, mice colonized with control *E. coli* showed higher infiltration of Ly6C^+^ and CCR2^+^ MPs in comparison to *Δ*(*atpA-atpG*) *E. coli*- colonized mice (**Fig 3B,C**). CSF2 governed the abundance of CCR2^+^ MPs and has previously been reported to be produced by ILC3s in a microbiota-dependent manner (Mortha *et al*., 2014; Satoh-Takayama *et al*., 2008). To determine the impact of ATP^ex^ on ILC3s, we analyzed CSF2 release and ILC3 counts in the colonic LP of untreated GF mice versus mice mono-colonized with *Δ*(*atpA-atpG*) or control *E. coli*. Notably, only mice gavaged with wild-type *E. coli* but not *Δ*(*atpA-atpG*) showed an increase in ILC3 numbers and CSF2 release, implicating the regulation of CSF2-producing ILC3s by microbe-derived ATP^ex^ (**Fig 3D**). In support of these data, decreased CSF2 production by ILC3s was observed in *Nlrp3*^-/-^ and *P2rx7*^-/-^ mice, deficient in initiating ATP^ex^-dependent inflammasome activation (**Fig 3E**). Altogether, these data indicate that microbe-derived ATP^ex^ regulates colonic monocyte-derived MPs and the production of CSF2 by ILC3s via the inflammasome.

### CSF2-producing ILC3s support intestinal CCR2^+^ Tim-4^-^CD4^-^ MPs

Microenvironmental cues within anatomical niches are critical for imprinting tissue-resident MP identity in various organs (Guilliams *et al*., 2020). However, less focus has been placed on niches for monocyte-derived MPs. In the gut, CSF2-producing ILC3s are abundantly found within postnatally formed tertiary lymphoid organs (TLOs), such as cryptopatches and isolated lymphoid follicles (Mortha *et al*., 2014). These data prompted us to determine whether CSF2-producing ILC3s constitute supporting cells for monocyte-derived MPs in the colon. In support of our hypothesis, live imaging of *Rorc*^+/EGFP^*Ccr2*^+/RFP^ colons revealed an accumulation of CCR2^+^ cells along the edges of TLOs, surrounding RORγt^+^ ILC3s within the structures (**Fig 4A**). To confirm that these CCR2^+^ cells surrounding TLOs are MPs, we quantified CX3CR1^+^CCR2^+^ MPs in the LP or TLOs of the colon in immunofluorescence images from *Cx3cr1*^+/EGFP^*Ccr2*^+/RFP^ mice. CCR2^+^ MPs displayed an elevated accumulation within TLOs compared to the surrounding LP (**Fig 4B,C**). TLOs have been reported to contain B cells and T cells (Hamada *et al*., 2002). To determine whether B and T cells were involved in shaping intestinal MP heterogeneity, MP composition was analyzed in WT, *Rag2^-/-^* and *Rag2^-/-^Il2rg^-/-^* mice. Interestingly, only *Rag2*^-/-^*Il2rg*^-/-^ mice (lacking all lymphocytes), but not *Rag2*^-/-^ mice (sufficient in ILCs), displayed a significant reduction in colonic Tim-4^-^CD4^-^ MPs that was comparable to the decrease observed in *Csf2*^-/-^ mice. These findings indicate that ILCs but not T and B cells regulate the homeostatic CSF2-dependent MP composition in the colon (**Fig 4D**). In line with this, adoptive transfer of 10^4^ FACS-purified *Rorc*^+/EGFP^ ILC3s from WT (ILC3^CSF2^) or *Csf2*^-/-^ (ILC3^ΔCSF2^) mice into *Rag2*^-/-^*Il2rg*^-/-^ recipients revealed a significant accumulation of Tim-4^-^CD4^-^ MPs in the colonic LP after 6 weeks (**Fig 4E, S4A**). The accumulation of Tim-4^-^CD4^-^ MPs was critically dependent on ILC3-derived CSF2 (**Fig 4E, S4B**). Notably, numbers of Tim-4^-^CD4^+^ and Tim-4^+^CD4^+^ MPs were also slightly increased in *Rag2^-/-^Il2rg^-/-^* mice after transfer of ILC3^CSF2^, suggesting a partial dependency of this subpopulation on CSF2-producing ILC3s (**Fig 4F**). In summary, these findings demonstrate that CCR2^+^ MPs accumulate around colonic TLOs and that colocalization with CSF2-producing ILC3s supports Tim-4^-^CD4^-^ MPs in the colon.

**Figure 4.**
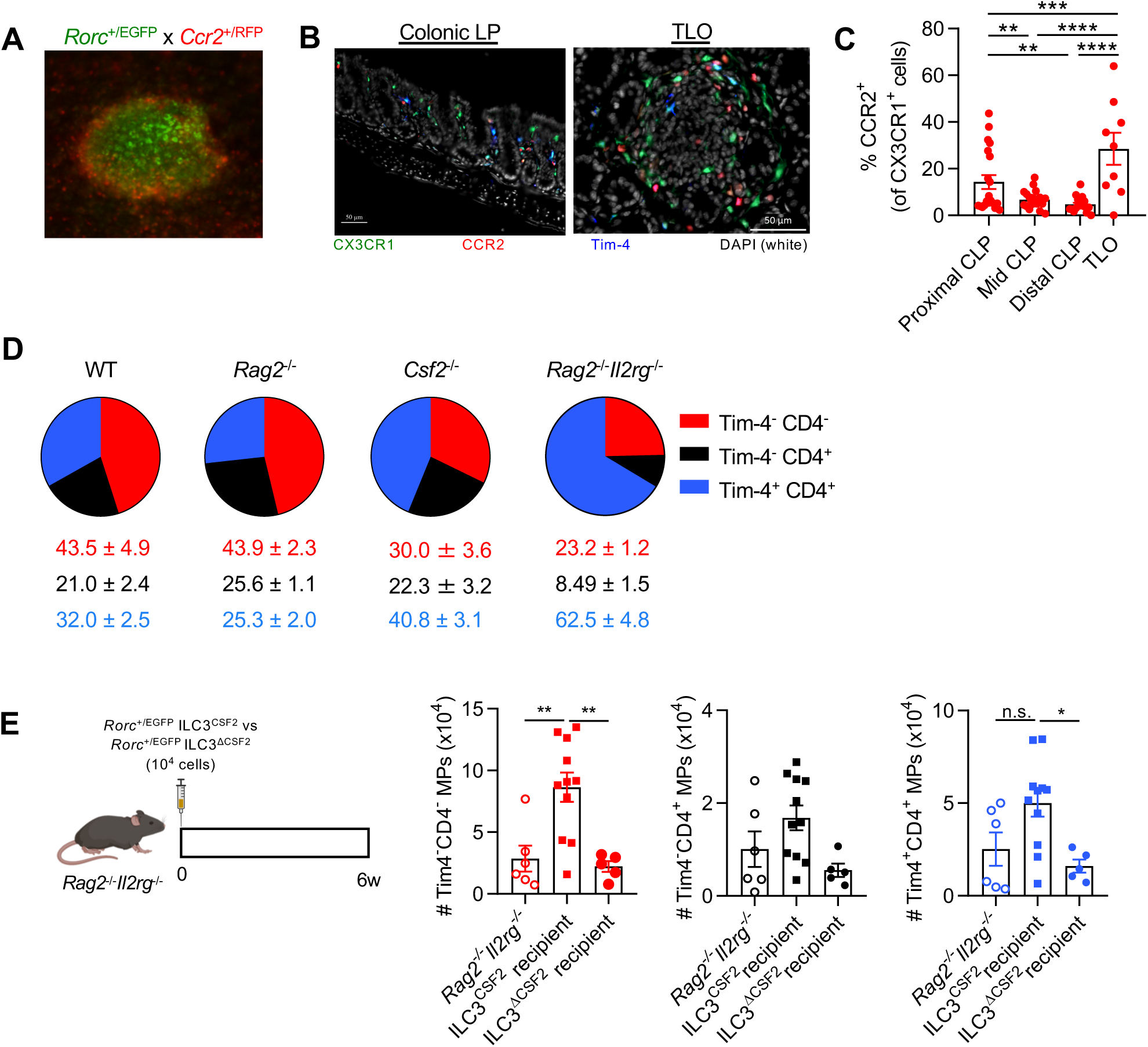
CSF2-producing ILC3s in TLOs induce CCR2^+^Tim-4^-^CD4^-^ MPs. (A) Representative live image of a colonic tertiary lymphoid organ (TLO) in *Rorc*^+/EGFP^*Ccr2*^+/RFP^ mice. (B) Representative immunofluorescence images of colonic lamina propria (LP) and TLOs of *Cx3cr1*^+/GFP^*Ccr2*^+/RFP^ mice stained for Tim-4 and DNA. (C) Proportion of CCR2-RFP^+^ of CX3CR1-GFP^+^ cells based on CellProfiler quantification of images in each colonic LP (CLP) region and TLOs, as indicated. (D) Colonic MP composition in adult sex-matched littermate mice as indicated. (E) 10^4^ FACS-sorted small intestinal ILC3s from either WT or *Csf2*^-/-^ mice were adoptively transferred into *Rag2*^-/-^*Il2r*^-/-^ mice, and recipients were analyzed at 6 weeks (left). Quantifications (right) of colonic MP populations post-transfer, as indicated. Data shown are representative of at least three independent experiments with at least three mice per group per experiment. One-way ANOVA with post-hoc Tukey’s test was performed for (C) and (E); *p < 0.05, **p < 0.01, ***p < 0.001, ****p < 0.0001; n.s., not significant.

### scRNA-Seq reveals unique TLO-associated MP populations

To incorporate spatial information into the actions of CSF2 on colonic MPs at higher granularity, we performed scRNA-Seq analysis of MPs isolated from either the TLOs or LP of WT or *Csf2*^-/-^ mice. Live *Cx3cr1*^+/EGFP^*Ccr2*^+/RFP^ colonic tissues revealed TLOs and LP using a fluorescence stereomicroscope (**Fig 5A**). Biopsy punches containing colonic TLOs or LP, free of TLOs, were isolated and digested prior to enrichment for CD11b^+^ cells by magnetic beads (>90% purity) and subsequent scRNA-Seq analysis (**Fig 5B**). UMAP dimensionality reduction and combined analysis of all 4 groups (LP^WT^, TLO^WT^, LP*^Csf2^*^-/-^, TLO*^Csf2^*^-/-^) yielded 18 clusters from a total of 15,369 cells (**Fig S4A**). Based on their top 30 cluster-defining genes, we identified clusters corresponding to B cells (clusters 7, 10), T/NK cells (cluster 14), and epithelial and stromal cells (clusters 0, 5, 6, 9, 11, 15-17), which were excluded from subsequent analysis (**Fig S4B**). DC clusters (3, 12, 13) were identified based on *Flt3*, *Dpp4*, *Zbtb46*, and *Itgax* expression. The remaining clusters (1, 2, 4, 8) were identified as MPs and monocytes based on their expression of *Csf1r*, *Cx3cr1*, and *C1qa* and absence of DC markers (**Fig S4C, D**).

**Figure 5.**
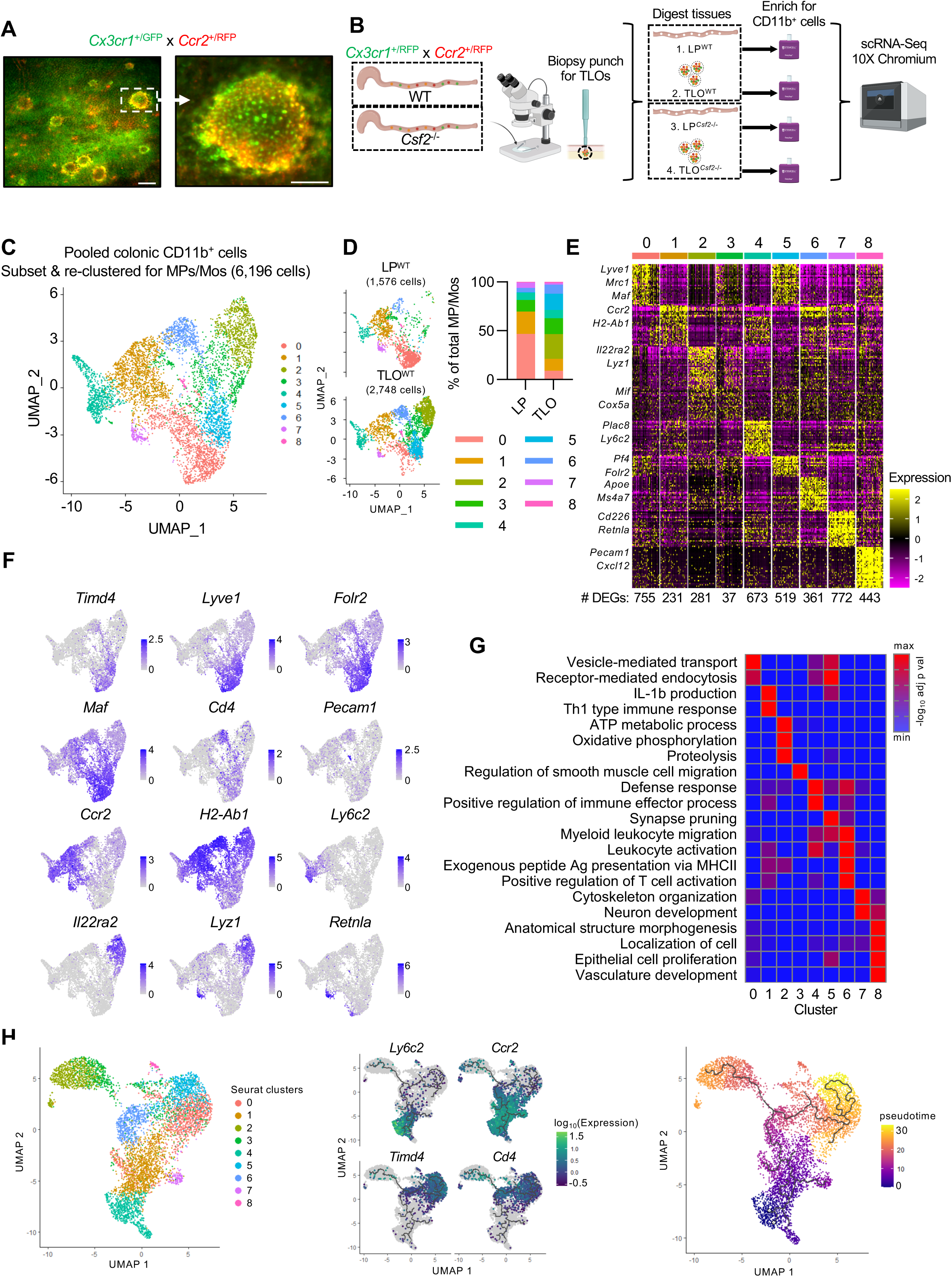
scRNA-Seq analysis of colonic LP MPs inside and outside of TLOs reveal further heterogeneity and preferential localizations. (A) Representative live image of a colonic tertiary lymphoid organ (TLO) in *Rorc*^+/EGFP^*Ccr2*^+/RFP^ mice. (B) Experimental scheme for scRNA-Seq set-up. (C) UMAP projection of the combined analysis of LP^WT^, TLO^WT^, LP*^Csf2^*^-/-^, and TLO*^Csf2^*^-/-^ subsetted and re-clustered for MPs and monocytes visualized together. (D) UMAP projection of MPs/Mos of LP^WT^ and TLO^WT^ (left) with quantification of the relative abundance of each cluster for each sample (right). (E) Heatmap depicting the top 30 DEGs for each cluster (log2FC threshold = 0.25, min.pct = 0.25, adjusted p <0.05), downsampled to 50 cells for visualization. The number of DEGs in each cluster is shown (bottom). (F) Feature plots illustrating expression of subset-defining genes. (G) Pathway enrichment analysis (gProfiler, Gene Ontology (GO) biological processes) using DEGs for each cluster. (H) UMAP dimensionality reduction using Monocle 3 was performed and visualized with overlaid Seurat annotations from (D) (left). Trajectory analysis was performed using Monocle 3 as indicated by solid black lines (middle, right). Changes in expression of subset-defining genes were visualized in conjunction with trajectory analysis (middle), and pseudotime analysis was performed and visualized using Monocle 3 (right).

Subsetting and re-analysis of the MP/monocyte clusters resulted in 9 distinct clusters, revealing substantial heterogeneity within the colonic MP pool (**Fig 5C**). Cells within clusters 0, 1, and 7 were enriched in the LP, while TLOs primarily comprised clusters 2, 3, 5, and 6, indicating preferential localization of some MP subpopulations within these structures (**Fig 5D**). Each MP cluster was then identified based on their top 30 cluster-defining genes (**Fig 5E**). MPs in clusters 0 and 5 highly expressed *Lyve1*, *Mrc1*, *Maf*, and *Timd4*, thus corresponding to tissue-resident Tim-4^+^ MPs (De Schepper *et al*., 2018; Dick *et al*., 2022; Moura Silva *et al*., 2021). MPs in clusters 0 and 5 co-expressed *Cd4* but not *Ccr2*, consistent with our flow cytometric classification (**Fig 5F**). Interestingly, MPs in cluster 5, enriched in the TLO, expressed higher levels of *Folr2*, which was shown to be expressed in gut and brain c-MAF-dependent perivascular MPs involved in metabolic regulation (Moura Silva *et al*., 2021). Consistent with previous reports, pathway analysis demonstrated that MPs in clusters 0 and 5 were enriched in endocytosis and vesicle-mediated transport pathways and mediated tissue homeostatic functions, including synapse pruning (**Fig 5G, Fig S4E**). These observations prompted us to label MP clusters 0 and 5 as Tim-4^+^ LP MPs and Tim-4^+^ TLO MPs, respectively. We also identified clusters corresponding to CCR2^+^ MPs (cluster 1), monocytes (cluster 4), Tim-4^-^CD4^+^ MPs (cluster 6), RELMα^+^ MPs (cluster 7), and epithelium/endothelium-associated *Pecam1*^+^ MPs (cluster 8) (**Fig 5E,F**). Although previous reports identified Tim-4^-^CD4^+^ MPs and investigated their developmental kinetics, the functions of this subpopulation remain unclear (Shaw *et al*., 2018). Interestingly, cluster 6 MPs, corresponding to Tim-4^-^CD4^+^ MPs, displayed a gene expression pattern similar to that reported in inflammatory microglia and border-associated MPs, including expression of *Apoe*, *Ms4a7*, and *Tmem119* (Amorim *et al*., 2022; De Schepper *et al*., 2018; Sankowski *et al*., 2019; Satoh *et al*., 2016). Pathway analysis on MP cluster 6 indicated that Tim-4^-^CD4^+^ MPs are involved in leukocyte activation, antigen presentation, T cell activation, and NF-κB signaling, suggesting a pro-inflammatory role for this subpopulation (**Fig 5G, S4E**).

Interestingly, cluster 2 and 3 MPs were found almost exclusively in TLOs and expressed high levels of *Il22ra2* and *Lyz1*, markers reported for TLO-residing CD11c^+^MHCII^+^CD11b^-^ CD103^-^ DCs in the small intestine (Guendel *et al*., 2020). However, we confirmed the MP identity for cluster 2 and 3 cells based on their expression of MP markers (*Csf1r*, *Cx3cr1*, *C1qa*, *Adgre1*, and *Fcgr1*) and the absence of DC markers (*Flt3*, *Dpp4*, and *Zbtb46*) (**Fig S4B,C**). The absence of *Timd4* and *Cd4* expression and low level of *Ccr2* expression in cluster 2 and 3 MPs suggest that these cells correspond to a subset of the Tim-4^-^CD4^-^ MPs (**Fig 5F**). These MPs were enriched in ATP metabolism, oxidative phosphorylation, and phagocytic pathways, indicative of a population high in energy demand (**Fig 5G, S4E**). To determine the developmental relationship between TLO-enriched and LP- enriched MP clusters, we performed trajectory analysis using Monocle 3 with Seurat-generated clusters overlaid, which revealed a common origin for all MPs within the monocyte cluster 4, confirming our fate-mapping and parabiosis studies (**Fig 5H**). Pseudotime analysis revealed a gradual loss of *Ly6c2* and *Ccr2* expression and gain of *Cd4* and *Timd4* expression as cells transition from monocytes towards differentiated MPs (**Fig 5H**). Importantly, Tim-4^-^CD4^+^ MPs represent a differentiation branching point, at which cells either transition into Tim-4^+^ MPs (clusters 0 and 4) or into TLO-associated MPs (first to cluster 3, then cluster 2) (**Fig 5H**). Given that Tim-4^-^CD4^+^ MPs were least affected by perturbations in the microbiota (**Fig 2**), their status as a defining branchpoint for tissue-resident MPs may be worth additional investigation in the future. In summary, our scRNA-Seq analyses reveal a novel TLO-associated Tim-4^-^CD4^-^ MP population, high in energy metabolism and possibly originating from Tim-4^-^CD4^+^ MPs along a distinct differentiation pathway.

### CSF2 is a spatial determinant of MP development and function in the colon

To determine whether TLO MPs may be regulated by CSF2, we first identified CSF2-responsive MP clusters based on their *Csf2ra* and *Csf2rb* expression. All MP clusters with the exception of *Timd4*^+^*Lyve1*^+^ MPs (clusters 0, 5, and 8) expressed detectable levels of *Csf2ra* and *Csf2rb* (**Fig 6A**). Comparison of MP cluster composition in WT and *Csf2*^-/-^ TLOs revealed a loss in the relative abundance of cluster 2 and 3 MPs in TLO*^Csf2^*^-/-^ (**Fig 6B**). Surprisingly, Tim-4^+^ LP MPs (cluster 0) were reduced in the LP^Csf2-/-^, even in the absence of *Csf2ra* and *Csf2rb* mRNAs (**Fig 6B**). We hypothesized that CSF2 deficiency renders colonic monocytes and CCR2^+^ MPs unable to differentiate and undergo apoptosis, based on CSF2’s role as a pro-survival factor for myeloid cells (Wan *et al*., 2013). An assessment of apoptosis in colonic MPs revealed an increase in apoptotic Ly6C^+^ and Tim-4^-^CD4^-^ MPs in *Csf2*^-/-^ mice, the latter corresponding to the loss of TLO MPs in TLO*^Csf2^*^-/-^ (**Fig S5**).

**Figure 6.**
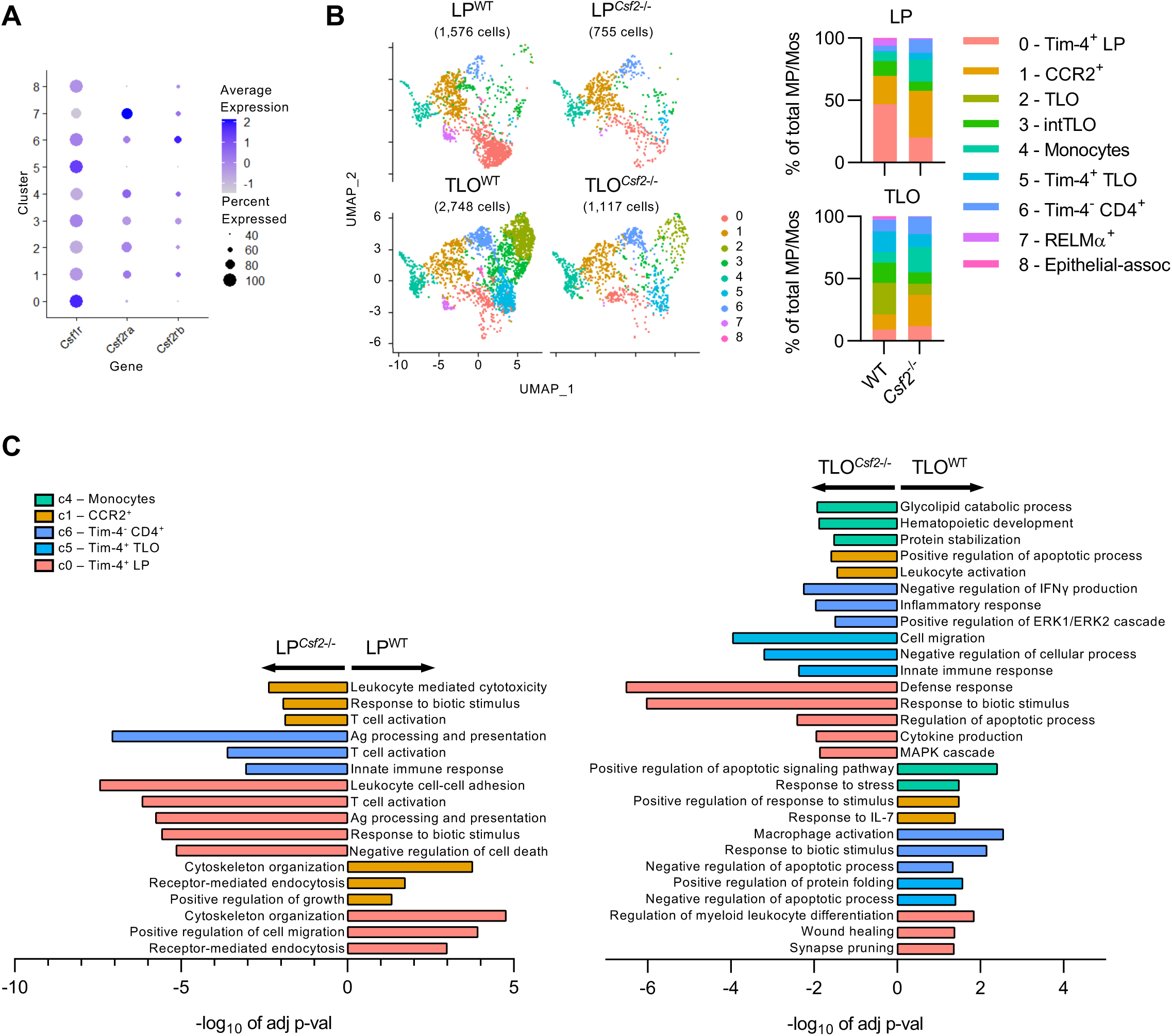
CSF2 deficiency results in a loss of TLO MPs and functional dysregulation of colonic MPs. (A) Dot plot showing expression of *Csf1r*, *Csf2ra*, and *Csf2rb* in each cluster from the merged data. Color denotes expression level, and dot size indicates percent of cells within the cluster expressing the gene, as indicated. (B) UMAP projection of MPs/Mos of each sample (left) with quantification of the relative abundance of each cluster for each sample (right). (C) Pathway enrichment analysis (gProfiler, Gene Ontology (GO) biological processes) using DEGs differentially regulated between WT versus *Csf2*^-/-^ in each region (LP, left; TLO, right) for selected clusters, as indicated. Negative values of -log_10_ of adjusted p-value indicate upregulation in *Csf2*^-/-^, positive values indicate upregulation in WT.

CSF2 promotes the transition of monocyte to MPs in the inflamed brain, specifically by supporting disease-promoting functions in MPs (Amorim *et al*., 2022). These new findings are in contrast to our data showing CSF2-driven pathways in steady state colonic MPs. Pathway analysis based on differential gene expression of each cluster in each region of WT versus *Csf2^-/-^* mice revealed significant differences in multiple MP clusters, even those devoid of *Csf2ra* or *Csf2rb* mRNAs (**Fig 6C**). Within the LP*^Csf2-/-^*, Tim-4^+^ (red) and CCR2^+^ (orange) MPs were impaired in cytoskeleton organization, migration, and endocytosis, but enriched in pro-inflammatory processes like T cell activation, antigen activation, leukocyte-mediated cytotoxicity, and the response to biotic stimuli (**Fig 6C**). However, an even larger number of MP clusters were affected in TLO*^Csf2-/-^*. Similar to LP*^Csf2-/-^* Tim-4^+^ MP, homeostasis of TLO*^Csf2-/-^* Tim-4^+^ (aqua) MPs was impaired as shown by altered synapse pruning and wound healing functions, and upregulated pro-inflammatory pathways (**Fig 6C**). Tim-4^-^CD4^+^ (dark blue) MPs followed these trends towards altered homeostasis and increased inflammation. Conversely, clusters corresponding to monocytes (green) and CCR2^+^ MPs downregulated pathways involved in response to stimuli and were enriched in pathways related to cell survival, glycolipid catabolism and protein stabilization in TLO*^Csf2-/-^*, confirming the increased apoptosis in *Csf2^-/-^* mice (**Fig S5A and 6C**). These data show that CSF2 deficiency induces the functional dysregulation of multiple colonic MP populations, particularly in the TLO. As a result, *Csf2^-/-^* colons are enriched in MPs skewed towards pro-inflammatory processes, while downregulating functions attributed to anti-microbial host defense and homeostasis. Collectively, these findings demonstrate that CSF2-rich TLOs constitute microanatomic niches for the homeostatic development and functional programming of MPs in the colon.

### CSF2-dependent Tim-4^-^CD4^-^ MPs are required for proper host defense against enteric infections

ILC3s are central for the innate immune response against attaching and effacing enteric pathogens like *Citrobacter rodentium*, the murine counterpart of human enteropathogenic *E. coli* (Song *et al*., 2015). CSF2 has been implicated in supporting anti-microbial host defense against *C. rodentium* through the modulation of CD11c^+^ myeloid cells (Hirata *et al*., 2010). Assessment of the colonic MP composition in WT and *Csf2*^-/-^ mice after infection with *C. rodentium* revealed a significantly altered expansion of Tim-4^-^CD4^-^ MPs in the absence of CSF2 (**Fig 7A**). In contrast to previous reports using *C. rodentium* infections, CD11c^+^MHCII^+^CD64^-^ DCs did not significantly differ in *Csf2^-/-^* and littermate controls post infection (**Fig S6A**). While differences in the distribution of cDC1 and cDC2 were observed, these results suggest that CSF2-dependent MPs mediate efficient anti-microbial host defense (**Fig S6B**). Consequently, *Csf2*^-/-^ mice showed greater weight loss, accompanied by higher bacterial burden and dissemination despite comparable infection efficiency (**Fig 7B and C**).

**Figure 7.**
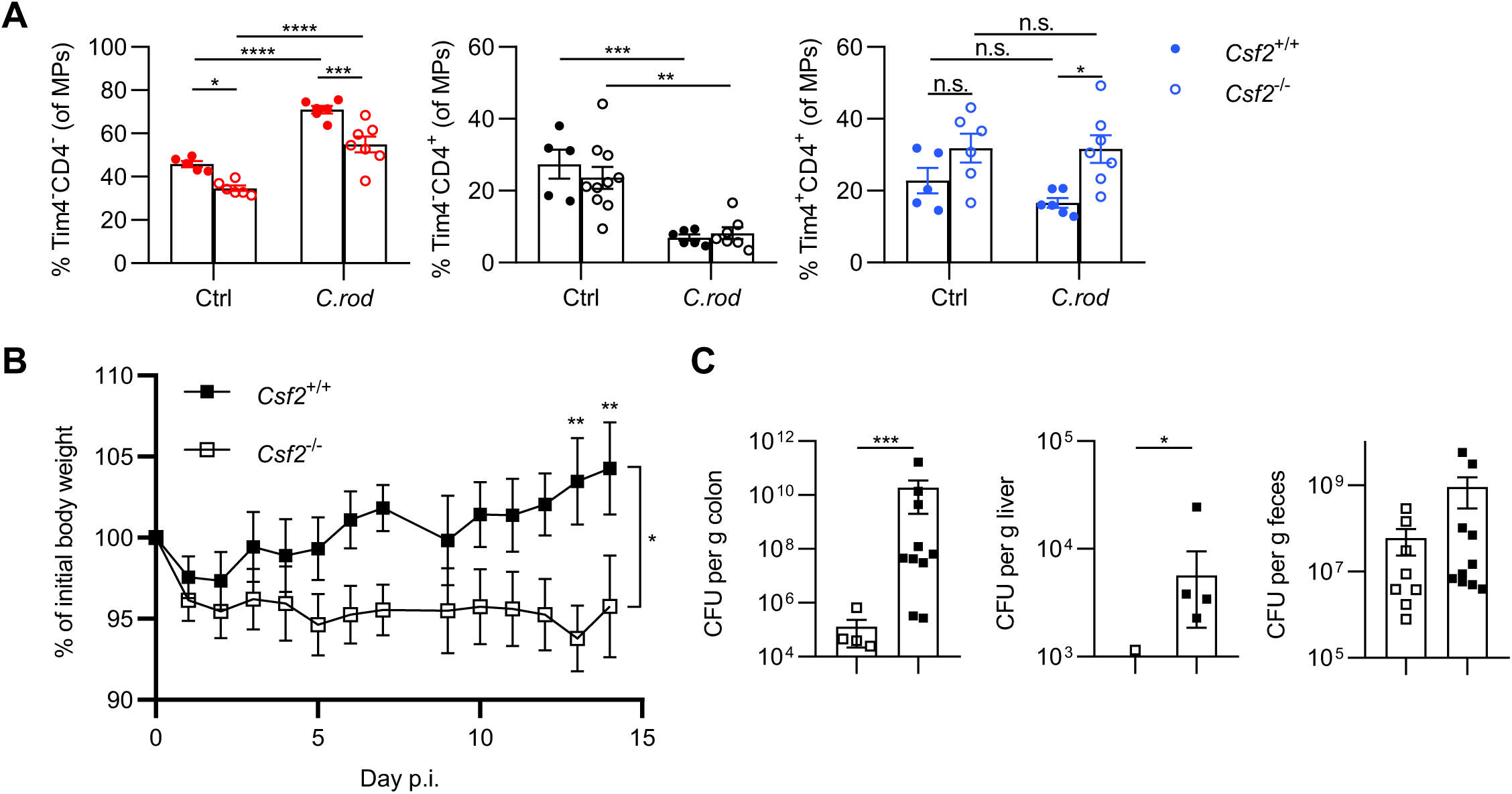
Host defense against *C. rodentium* requires CSF2-dependent Tim-4^-^CD4^-^ MPs. Groups of age- and sex-matched mice were infected with *C. rodentium* for 2 weeks. (A) Quantification of relative abundance of colonic MP populations. (B) Body weight was tracked daily during the course of infection. (C) Quantification of *C. rodentium* in the colon, feces, and liver at end point. Data shown is representative of at least three independent experiments with at least three mice per group per experiment. Two-way ANOVA with post-hoc Tukey’s multiple comparison test was performed for (A), mixed-effects analysis was performed with post-hoc Sidak’s multiple comparisons test for (B), and unpaired Student’s t test was performed for (C); *p < 0.05, **p < 0.01, ***p < 0.001; n.s., not significant.

## Discussion

Intestinal MPs are critical for gut homeostasis and host defense. Uncovering the mechanisms regulating their developmental and functional heterogeneity across the largest mucosal surface is a critical yet challenging step towards a detailed understanding of these cells during steady state and inflammation. Here, we provide new insights into the molecular and cellular interactions that govern microbiota- and host-regulated MP development and function in the colon. We demonstrate that microbe-derived ATP^ex^ fuels NLRP3-driven production of CSF2 by ILC3s. We identify TLOs as microanatomic locations for the interaction of ILC3s and monocyte-derived, CSF2-dependent MPs, representing a distinct niche for myeloid development and functional programming in the gut to support enteric anti-microbial host defense.

Intestinal TLOs form postnatally, or in response to chronic inflammation and colorectal cancer in a microbiota-dependent manner (Eberl and Lochner, 2009; Koscso et al., 2020; Overacre-Delgoffe et al., 2021). Although best known for their role in T cell-independent B cell maturation, our findings indicate that TLOs also serve as an activation hub for monocyte-derived MPs (Tsuji *et al*., 2008). While a population of TLO-located *Cxcl13*-expressing MPs support IgA-producing B cells during *Salmonella* infection, their developmental origin remains unknown (Koscso *et al*., 2020). We show that monocytes transition into TLO-associated, CSF2-dependent MPs to support host defense against infections. Whether monocyte-derived, TLO-associated MPs regulate adaptive immunity within TLOs in the steady state or inflammation remains an intriguing question for future investigation.

Collectively, our findings extend our knowledge on how the microbiota contributes to the steady state heterogeneity of intestinal MPs. In contrast to bacteria-derived metabolites like tryptophan or short-chain fatty acids, which require specialized biochemical pathways not present in all microbes, ATP is abundantly produced across all microbial kingdoms and governs the steady state activation of ILC3s. Our findings indicate that ATP^ex^, as a measurement for universal microbial energy metabolism, may serve as a rheostat for the control of gut MP development. The instability of ATP and expression of ecto-nucleotidases by the intestinal epithelium may be a rate limiting step in the activation of ILC3s and the regulation of MP heterogeneity, opening new avenues for exploration into these interactions.

While ATP^ex^-mediated activation of the inflammasome exacerbates disease in extra-intestinal locations, it constitutes a critical element to tune immune homeostasis (Aganna *et al*., 2002). In contrast to its pro-inflammatory role in other organs, CSF2 governs the steady-state transition of monocytes to macrophages in the colon and controls essential homeostatic functions and defense pathways across multiple gut MP subpopulations (Achuthan et al., 2021). ILC3-derived CSF2 promotes survival of monocytes and shapes the metabolic program of colonic monocyte-derived MPs in TLOs, while maintaining functional specification of other gut-resident MP subpopulations. Moreover, TLO- associated MPs follow a distinct developmental trajectory compared to LP MPs diverging from Tim-4^-^CD4^+^ MPs. Deficiency in *Csf2* results in the loss of TLO-associated MPs and promotes inflammatory pathways in LP MPs even if their gene expression suggests unresponsiveness to CSF2. This enrichment of inflammatory pathways in *Csf2*-deficient MPs may pose as a coping strategy to prevent infections and is of translational relevance, considering the presence of mutations in *CSF2RB* and the presence of neutralizing anti-CSF2 autoantibodies in complicated Crohn’s disease (CD) (Chuang *et al*., 2016; Han *et al*., 2009). Interestingly, neutralizing anti-human CSF2 autoantibodies precede the onset of CD by several years (Mortha, 2021). This suggests that such perturbations of the steady state, microbiota-triggered, ILC3-CSF2 niche for MP development and function may raise the susceptibility of enteric infections that may possibly contribute the onset of CD (Mortha, 2021).

Collectively, our data identify a previously underappreciated role for microbe-derived ATP as a regulator of a CSF2-dependent tissue niche for the development of TLO-residing monocyte-derived MPs that support host defense in the healthy intestine.

## Acknowledgments

We thank all members of #theonlylabever for their support and discussion. We acknowledge support by the University of Toronto, Temerty Faculty of Medicine Flow Cytometry Core facility, the 10x Genomics staff at the Princess Margaret Genomics Centre, and the Division for Comparative Medicine. We wish to thank Dr. Juan-Carlos Zúñiga-Pflücker and Dr. Adam Gehring for critical reading of the manuscript.

This study was supported by an Ontario Trillium Scholarship and Vanier Canada Graduate Scholarship - NSERC (P.C.). K.B. is supported by a Canadian Institutes of Health Research (CIHR) Banting Postdoctoral Fellowship Program and L.N. by an Ontario Graduate Scholarship and a NSERC-PGS award. S.L.T. is a recipient of the Dr. Edward Ketchum Graduate Student Scholarship and the Canada Graduate Scholarships – Master’s (CGS M) award. T.M. is supported by a Canada Research Chair in NKT cell Immunobiology, a CIHR grant (PJT-175055), and a Canada Foundation for Innovation Physical Infrastructure Grant (29186). A.M. is supported by the Canadian Foundation for Innovation John R. Evans Leaders Fund, the CIHR (PJT-388337, 6210100847, 6210100850) and a NSERC-Discovery Grant (RGPIN-2019-04521). A.M. is the Tier 2 Canadian Research Chair in Mucosal Immunology and supported by the Tier 2 CRC-CIHR program (CRC-2021-00511).

## Author contributions

Conceptualization, P.C. and A.M.; Methodology, P.C. and A.M.; Software, P.C. and H.H.; Investigation, P.C., K.B., L.N., E.Y.C., S.L.T., H.L., A.W., M.K., T.D., Ab.Mo., and S.M.L.; Writing – Original Draft, P.C. and A.M.; Writing – Review & Editing, P.C. and A.M.; Funding Acquisition, A.M.; Resources, T.M., T.C., H.I., and S.E.; Supervision, A.M.

## Figure legends

**Supplemental Figure 1.**
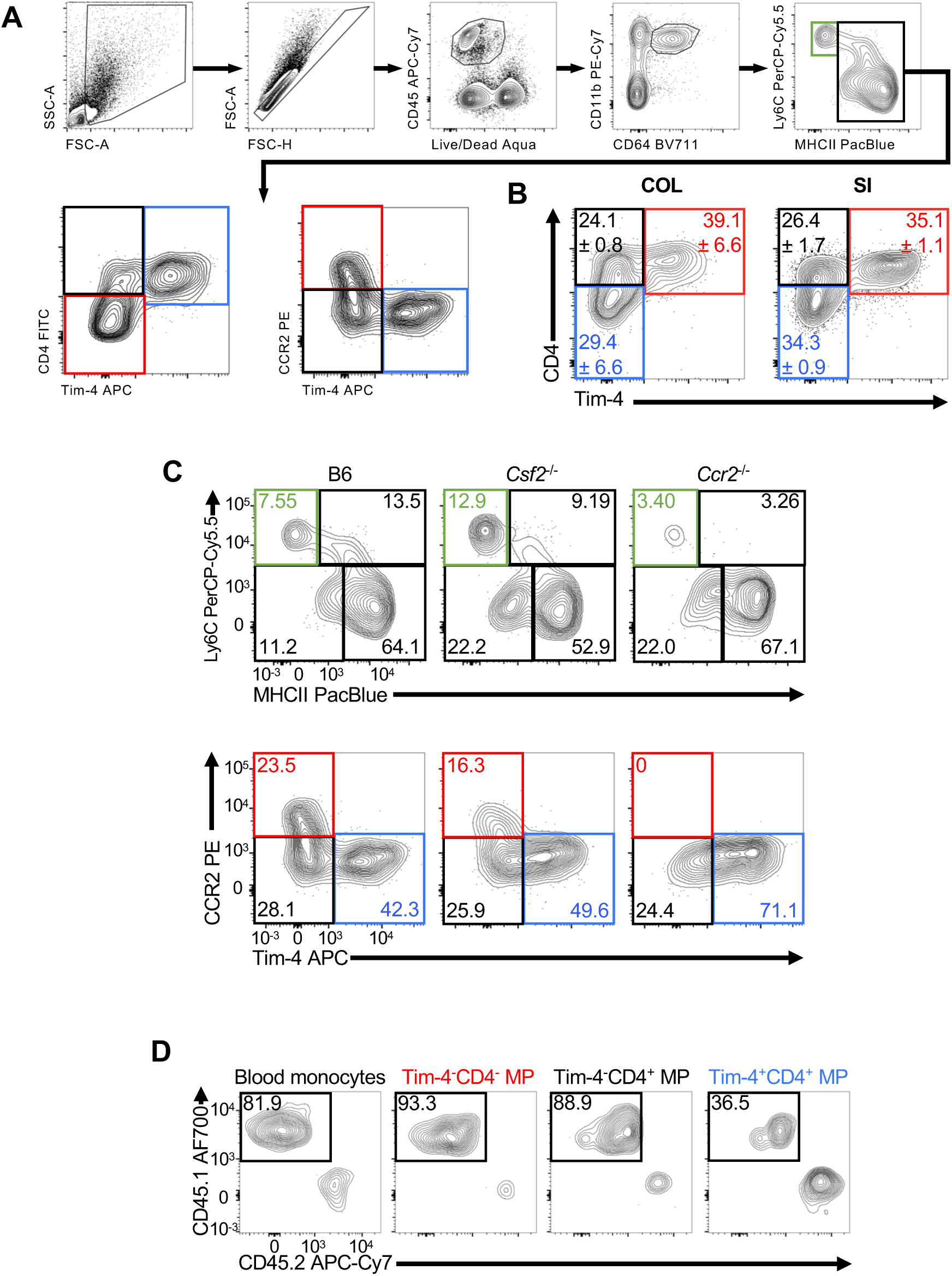
Macrophage classification and developmental phenotyping. (A) Gating strategy for intestinal MPs. (B) Representative flow cytometry plots of colonic and small intestinal MPs using the Tim-4/CD4 classification. (C) Representative flow cytometry plots of colonic MPs in sex-matched littermate WT, *Csf2*^-/-^, and *Ccr2*^-/-^ mice showing the monocyte waterfall (top) and Tim-4/CCR2 (bottom) gating strategies. (D) Representative flow cytometry plots showing CD45.1 chimerism of colonic MP populations in 6-month parabiotic mice.

**Supplemental Figure 2.**
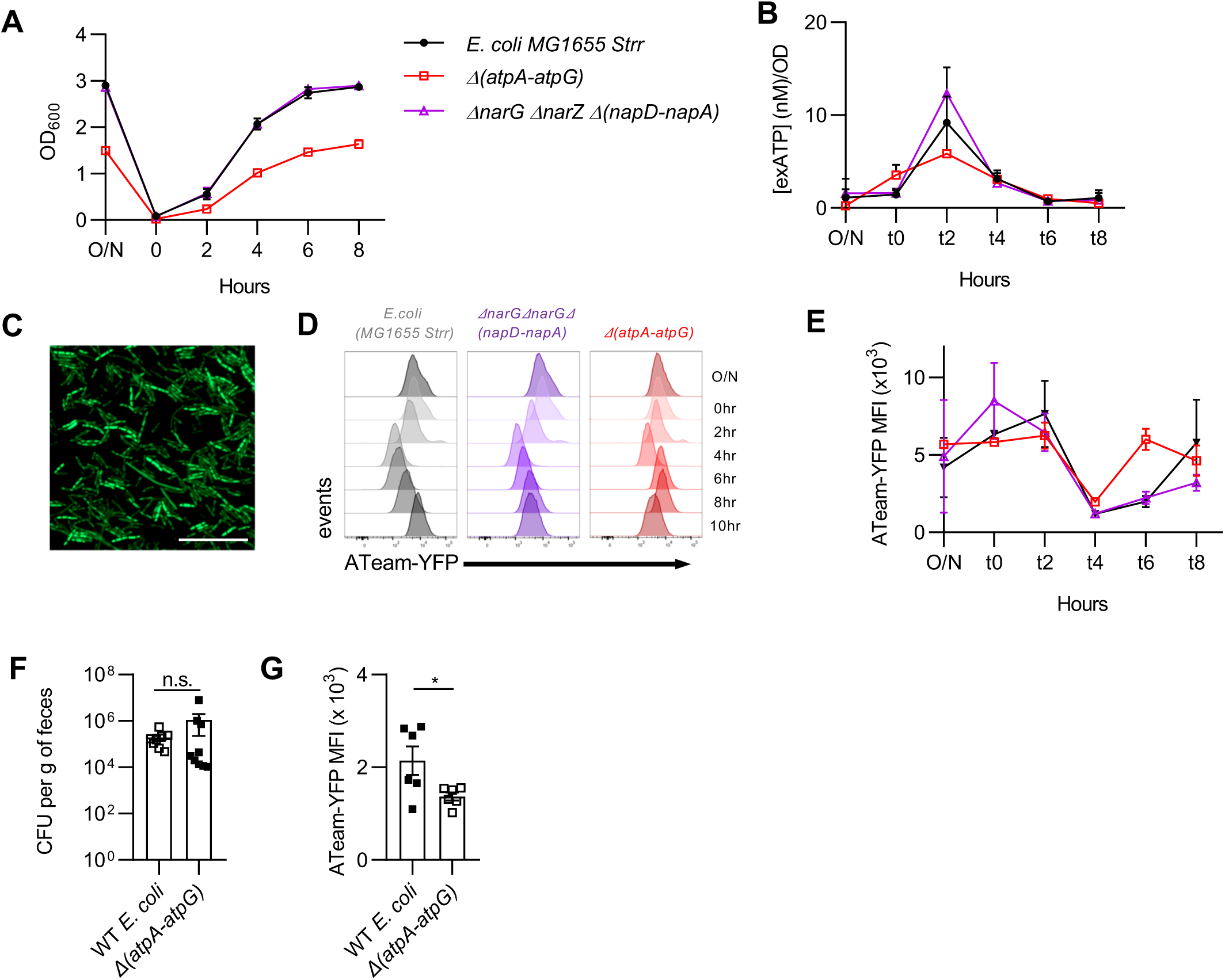
ATPase-deficient *E. coli* generate lower levels of ATP. (A-. E) *E. coli* MG1655 wild-type, ATPase-deficient (*Δ(atpA-atpG)*), and nitrate reductase-deficient (*ΔnarG ΔnarZ Δ(napD-napA)*) mutant strains were cultured *in vitro* overnight, then transferred into fresh media, and subsequently evaluated at the indicated timepoints. (A) Growth curve of *E. coli* MG1655 strains measured by OD_600_ as indicated. (B) Supernatant ATP levels (ATP^ex^) were quantified using the Promega ENLITEN ATP Assay as per the manufacturer’s instructions. (C) Intracellular ATP (ATP^int^) can be quantified by measuring fluorescence intensity of the ATeam 3.10 plasmid. (D) Representative histograms showing fluorescence intensity of the ATeam 3.10 plasmid in each *E. coli* strain at the specified timepoints measured by flow cytometry. (E) Quantification of the median fluorescence intensity of (D). (F-G) *E. coli* MG1655 wild-type and *Δ(atpA-atpG)* mutant were isolated and assessed from 1-week-colonized germ-free mice. (F) CFUs of each *E. coli* strain were quantified in the feces of colonized mice. (G) Each *E. coli* strain was isolated from fecal samples of respective colonized mice and assessed for levels of ATP^int^ by flow cytometry; showing median fluorescence intensity (MFI). Data shown are representative of at least three independent experiments with at least three mice per group per experiment. Unpaired Student’s t test was performed for (F) and (G); *p < 0.05; n.s., not significant.

**Supplemental Figure 3.**
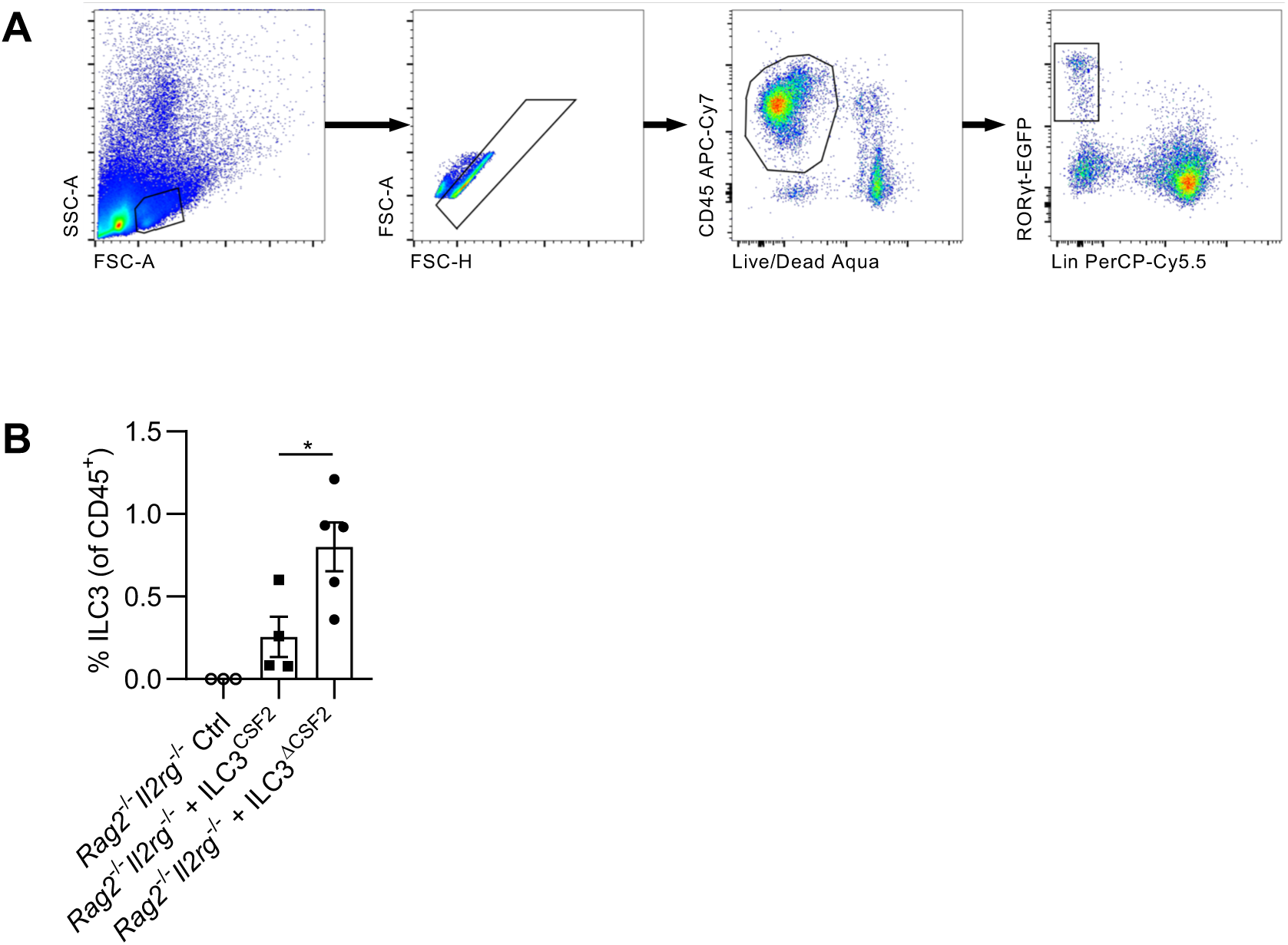
Sorting strategy and post-transfer verification of ILC3s for adoptive transfer experiment. (A) Representative sorting strategy of SI *Rorc*^+/EGFP^ ILC3s (WT or *Csf2*^-/-^) for adoptive transfer. (B) Relative abundance of colonic ILC3s (Lin^-^ ROR*γ*t-GFP^+^) out of CD45^+^ cells to validate the reconstitution of *Rorc*^+/EGFP^ ILC3s in *Rag2*^-/-^*Il2r*^-/-^ recipients at 6 weeks post-transfer. One-way ANOVA with Tukey’s test was performed; *p < 0.05.

**Supplemental Figure 4.**
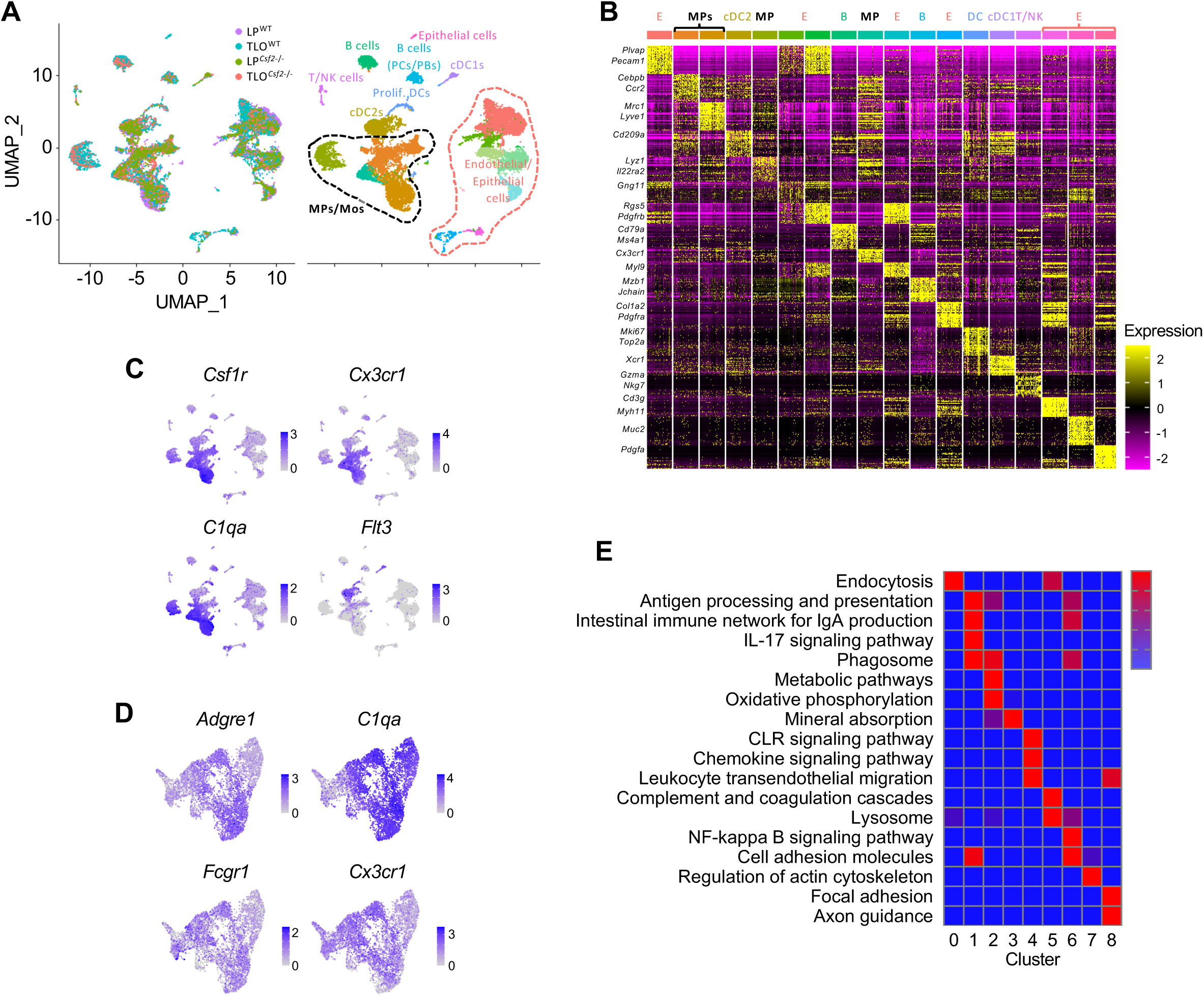
Gene expression profiling of macrophage clusters by scRNA-Seq. (A) UMAP dimensionality reduction and combined analysis of all datasets, representing 15,369 cells that have passed QC filtering; colored based on sample identity (left) or clusters (right). Cell populations identified and annotated based on DEG expression analysis of each cluster. (B) Heatmap of the top 30 genes per cluster, downsampled to 50 cells per cluster for visualization, based on DEG analysis across clusters (logFC threshold = 0.25, min.pct = 0.25, adj p val < 0.05). (C) Feature plots depicting gene expression patterns of MP/DC markers used to subset MP/Mo clusters and exclude DC clusters. (D) Feature plots confirming expression of MP markers in subsetted and re-clustered MP/Mos. (E) KEGG pathway enrichment analysis (gProfiler) using DEGs for each cluster.

**Supplemental Figure 5.**
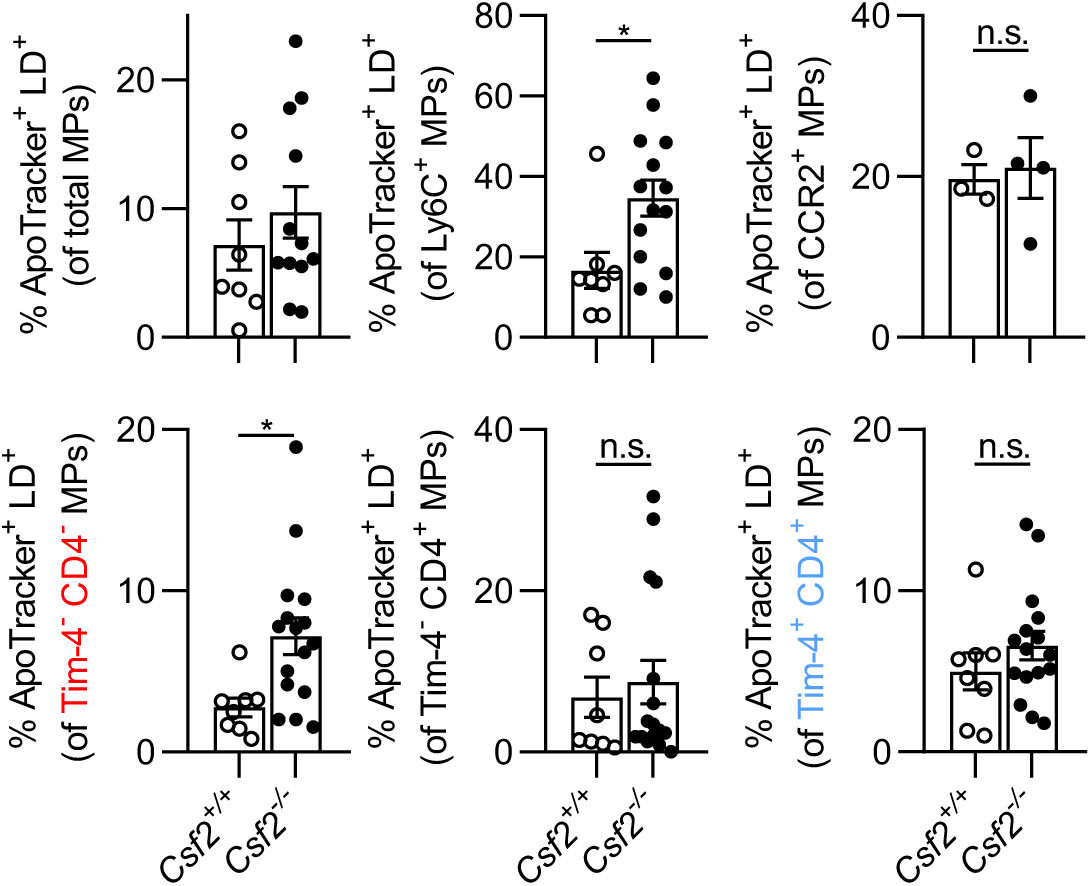
Differential analysis of WT versus *Csf2*^-/-^ MP subpopulations. Quantification of apoptotic cells within each colonic MP population in sex-matched littermate WT versus *Csf2*^-/-^ mice by ApoTracker staining, as indicated. Unpaired Student’s t test was performed; *p < 0.05; n.s., not significant.

**Supplemental Figure 6.**
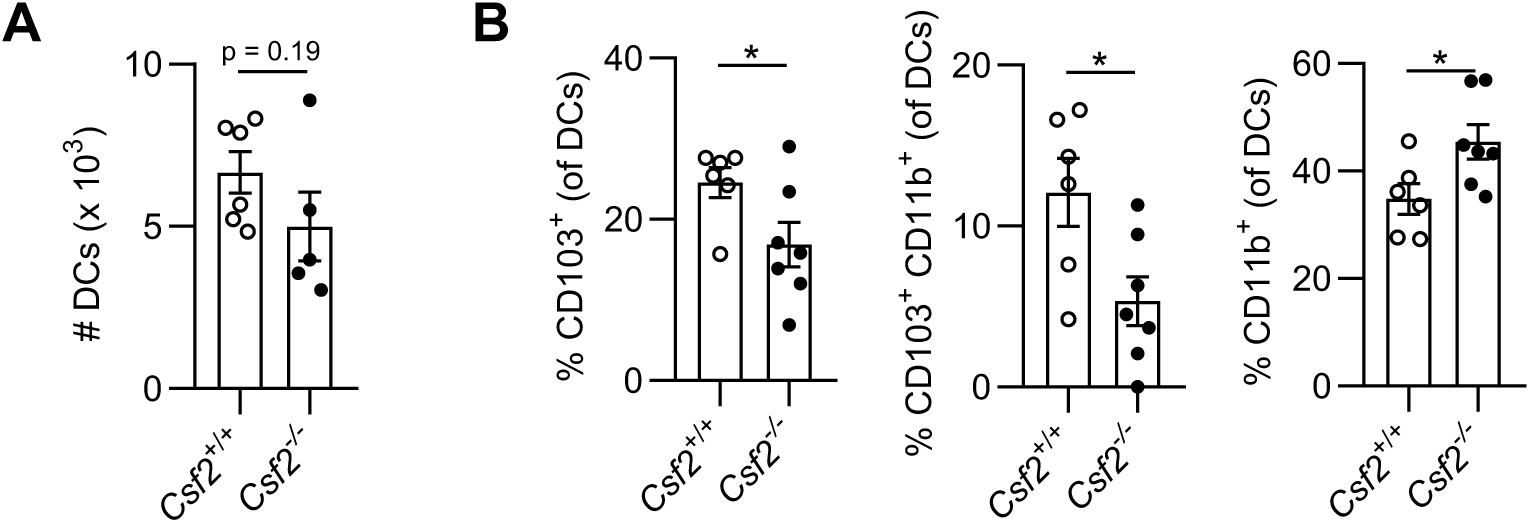
DC phenotyping during *C. rodentium* infection. Groups of age- and sex-matched mice were infected with *C. rodentium* for 2 weeks. (A) Quantification of CD11c^+^MHCII^+^CD64^-^ DCs. (B) Relative abundance of each DC subset. Unpaired Student’s t test was performed; *p < 0.05.

## STAR Methods

### Key resources table

For surface staining, the following anti-mouse Abs were used: TCR*β* (H57-597; eBioscience), CD4 (GK1.5; BioLegend), CD45 (30-F11; BioLegend), CD218a (IL-18Ra) (P3TUNYA; eBioscience), ST2 (RMST2-2; eBioscience), CD11b (M1/70; BioLegend), Ly6c (HK1.4; eBioscience), CD64 (X54-5/7.1; BioLegend), and MHCII (I-A/I-E) (M5/114.15.2; eBioscience). Intracellular markers include anti-mouse IFN-*γ* (XMG1.2; eBioscience), TNF*α* (MP6-XT22; eBioscience), IL-10 (JESS-16E3; BioLegend), IL-17A (TC11-18H10.1; BioLegend), and FOXP3 (MF-14; BioLegend). CD4^+^ T cells were gated as Live CD45^+^ TCRβ^+^ CD4^+^. Immature macrophages were gated as Live CD45^+^ CD64^+^ CD11b^+^ Ly6c^hi^ MHCII^lo^.

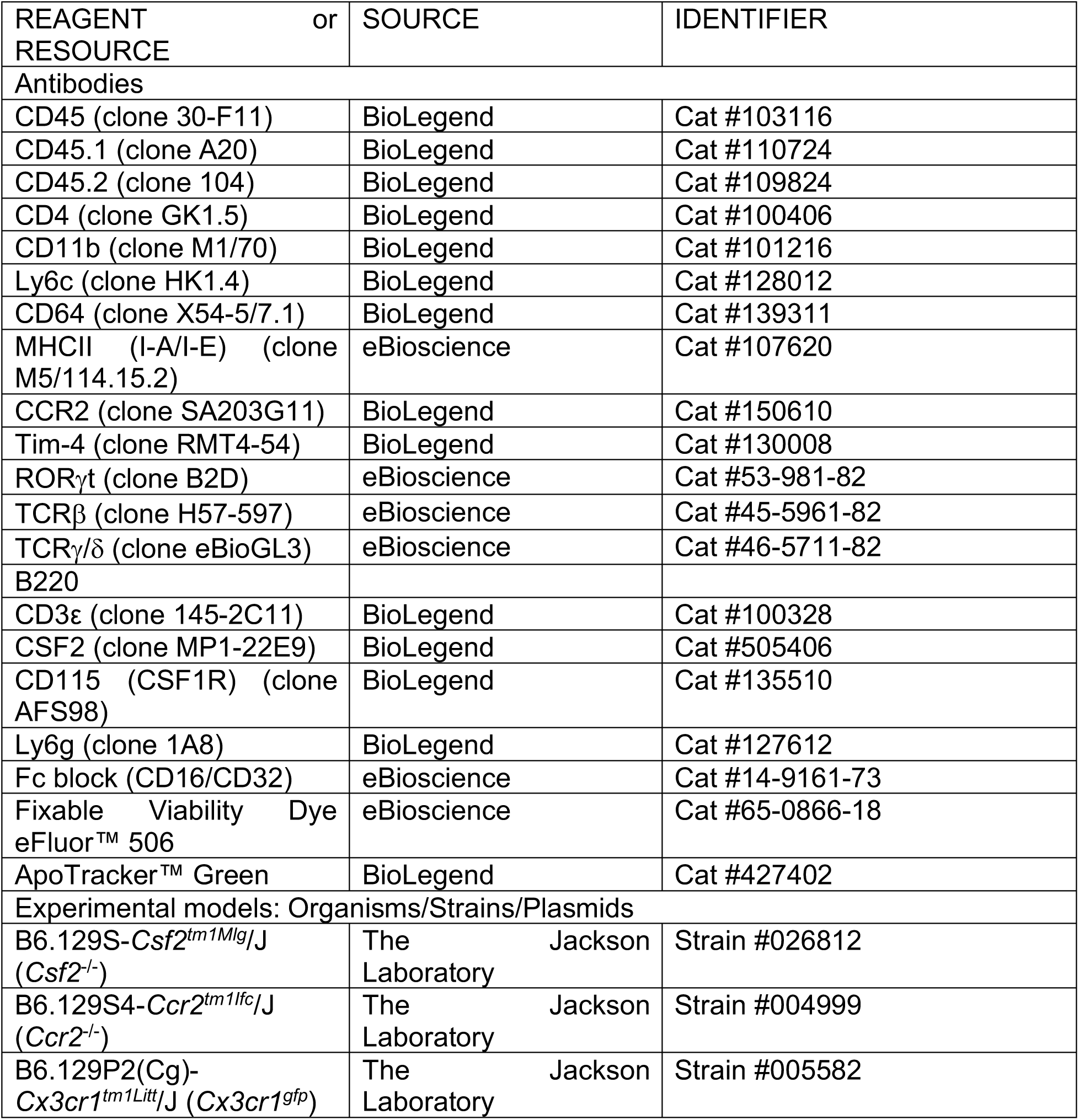
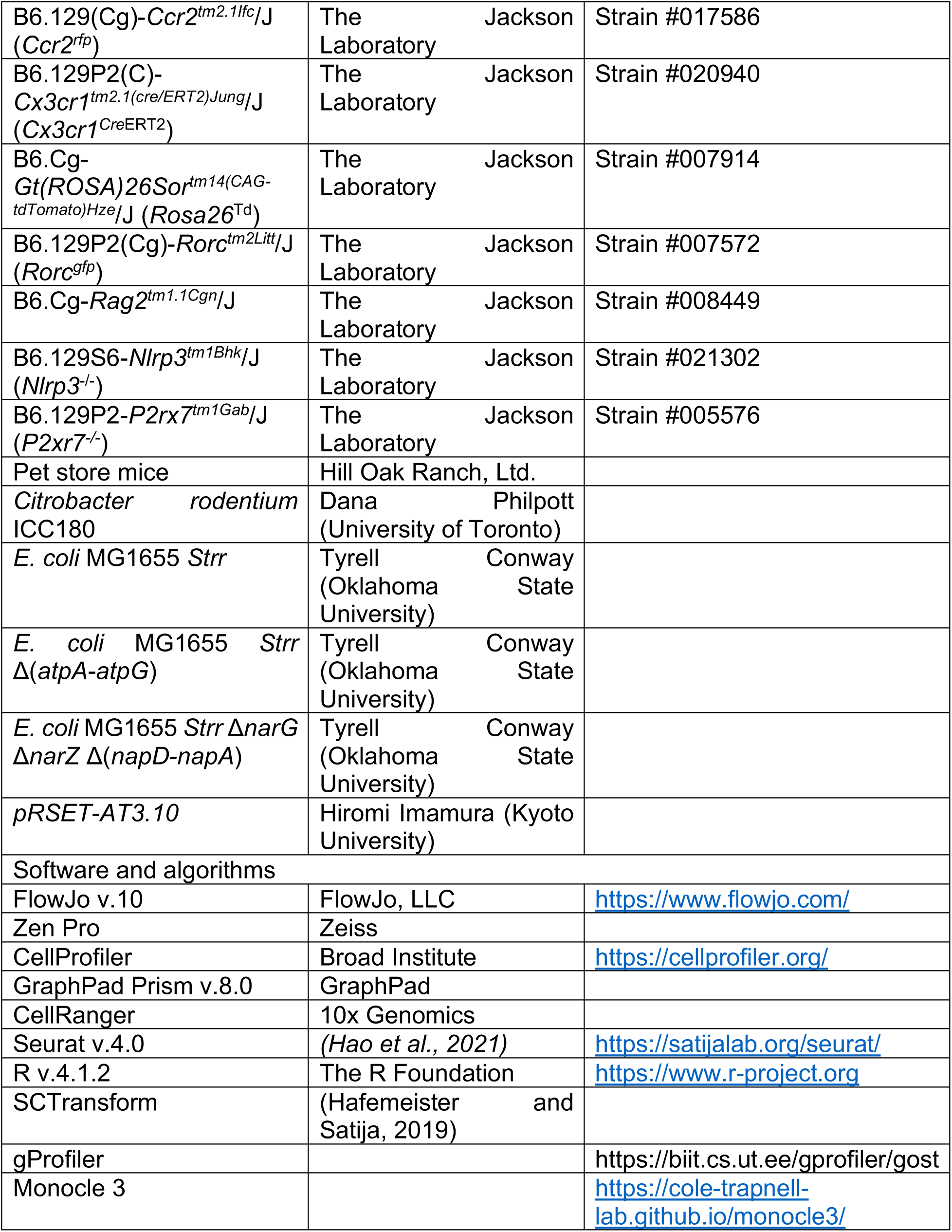

### Resource availability

Single-cell RNA-seq data additional information required to reanalyze the data reported in this paper is available from the lead contact upon request.

### Experimental model and subject details

#### Mice

All mice were purchased from Jackson Laboratory and subsequently bred in-house under specific pathogen-free conditions at the University of Toronto, Division of Comparative Medicine. Strains and stock numbers are listed in the Key resources table. Unless otherwise stated, all experiments were conducted using 8-10-week-old age- and sex-matched littermates. Germ-free animals were maintained in the gnotobiotic facility at the University of Toronto, Division of Comparative Medicine. To obtain re-wilded mice, pet store mice were purchased from High Oak Ranch Ltd. (Baden, ON) and bred in our mouse facility in a containment room (bioBUBBLE Inc, Fort Collins, CO). C57Bl/6 (B6) pups were co-housed with pet store pups from 3 to 7 weeks of age, separated and subsequently bred. B6 pregnant dams were gavaged with cecal content from pet store female mice 2 to 3 days prior to delivery. The pups were used to establish a re-wilded colony for experiments. All experiments were approved by the Faculty of Medicine and Pharmacy Animal Care Committee at the University of Toronto (animal use protocols 20011887 and 20012454 to TM and 20012400 to AM).

#### Microbes

All *E. coli* MG1655 strains were provided by Dr. Tyrell Conway (Oklahoma State University). *C. rodentium* ICC180 was a gift from Dr. Dana Philpott (University of Toronto). The plasmid encoding the ATP biosensors (pRSET-AT3.10) were a kind gift from Dr. Hiromi Imamura (Kyoto University).

### Method details

#### Purification and colonization of *Tritrichomonas musculis*

Cecal contents of *T. mu*^+^ mice were collected, resuspended in PBS, filtered through a 70 μm cell strainer, and spun for 10 min at 600 x *g*. The resulting pellet was put through a 40/80% Percoll gradient centrifugation. The *T. mu*-enriched interphase was collected. Protozoa were then resuspended in PBS and double sorted into PBS based on size, granularity, and violet autofluorescence on a FACSAria II. Two million *T. mu* were orally gavaged into mice immediately after the sort.

#### *C. rodentium* infection and pathological assessment

Groups of age- and sex-matched littermates were infected with *C. rodentium* ICC180 (∼2 x 10^8^ CFUs) by oral gavage as previously described (Bouladoux et al., 2017). Mice were weighed daily to monitor disease progression and euthanized at 2 w p.i. Colons were harvested for lamina propria leukocyte isolation and downstream analysis. Colony forming units (CFUs) of *C. rodentium* in feces, colon, and liver were measured on MacConkey agar plates containing 100 μg/mL kanamycin.

#### Generation of *E. coli* MG1655 ATeam strains

ATeam plasmid was isolated from *E. coli* ATeam3.10 using the Monarch^®^ Plasmid DNA Miniprep Kit (New England Biolabs) as per the manufacturer’s protocol. *E. coli* MG1655 *Strr, E. coli* MG1655 *Strr* Δ(*atpA-atpG*), and *E. coli* MG1655 *Strr* Δ*narG* Δ*narZ* Δ(*napD-napA*) strains were treated with calcium chloride to make them chemically competent for plasmid DNA uptake. Transformation was performed on these chemically competent cells to transfer the ATeam plasmid, following New England Biolabs’ High Efficiency Transformation Protocol.

#### Colonization of germ-free mice with *E. coli* strains

Groups of age- and sex-matched littermate germ-free mice were orally gavaged with ∼10^3^ CFU of *E. coli* MG1655 WT or *E. coli* MG1655 Δ*narG* Δ*narZ* Δ(*napD-napA*), or ∼10^4^ CFU of *E. coli* MG1655 *Strr* Δ(*atpA-atpG*). Differences in starting CFU accounted for the slower growth rate of *E. coli* Δ(*atpA-atpG*) to ensure equal colonization efficiency at time of analysis. Mice were analyzed 1 w later. Fecal pellets were collected both prior to gavage and at time of harvest to confirm colonization.

#### Antibiotics treatment

Mice were treated with metronidazole (0.5 g/L), ampicillin (1 g/L), neomycin (1 g/L), and streptomycin (1 g/L) *ad libitum* for 2 weeks via drinking water. Water containing antibiotics was exchanged every 3 days.

#### Isolation of intestinal lamina propria leukocytes

Colonic or small intestinal (SI) lamina propria (LP) cells were isolated as previously described (Chiaranunt et al., 2020). Briefly, intestines were washed in HBSS plus 5 mM EDTA and 10 mM HEPES to strip the epithelium. Tissues were then minced and shaken at 37°C for 20 min in digestion buffer (HBSS with calcium and magnesium, supplemented with 10 mM HEPES, 4% FBS, penicillin-streptomycin (Sigma Aldrich), 0.5 mg/mL DNase I (Sigma Aldrich), and 0.5 mg/mL Collagenase (Sigma Aldrich)). Supernatants were collected and enriched for leukocytes using a 40/80% Percoll gradient, after which cells are ready for downstream use.

#### Flow cytometry

For surface staining, after isolation of intestinal LP leukocytes, cells were resuspended in FACS buffer (PBS w/o Ca^2+^ Mg^2+^ supplemented with 2% heat inactivated FBS and 5 mM EDTA) and then incubated on ice for 20 min with Fc block (CD16/CD32; eBioscience), surface marker antibodies, and Fixable Viability Dye eFluor™ 506 (eBioscience). For flow cytometric detection of apoptotic cells, ApoTracker™ Green was added in conjunction to surface stains, and samples were instead incubated at room temperature for 20 min as per manufacturer’s protocol.

For intracellular staining, cells were first stimulated for 4 h at 37°C in R-10+ media supplemented with protein transport inhibitor cocktail containing Brefeldin A and Monensin (eBioscience). Cells were then washed and resuspended in FACS buffer and incubated on ice for 20 min with Fc block (CD16/CD32; eBioscience), surface marker antibodies, and Fixable Viability Dye eFluor™ 506 (eBioscience). Cells were fixed and permeabilized using the BD Cytofix/Cytoperm Kit, followed by cytokine stains, then re-fixed and permeabilized using the eBioscience Foxp3/Transcription Factor Staining Buffer Set, followed by transcription factor stains.

Samples were analyzed on an LSR Fortessa X-20 (BD) with subsequent cytometric data analysis using FlowJo. All antibodies used in this study are listed in the Key resources table.

#### *In vitro E. coli* culture and ATP measurement

Each *E. coli* strain was grown overnight at 37°C with shaking in LB broth containing 50 μg/mL streptomycin. The next day, OD600 was measured for each culture, and aliquots were taken for extracellular ATP (ATP^ex^) and intracellular ATP (ATP^int^) quantification. Each sample was then aliquoted into fresh media (LB with 50 μg/mL streptomycin) and placed in the shaking incubator. An aliquot was removed every 2 h for OD600 measurement and ATP quantification. For ATP measurements, aliquots were spun down at 3,000 x *g*. Supernatants were analyzed for ATP^ex^ using the ENLITEN ATP Assay System Bioluminescence Detection Kit (Promega) according to the manufacturer’s instructions. Pellets containing *E. coli* were resuspended and analyzed on the BD LSRFortessa for ATP^int^.

#### Luminal ATP measurement

Fecal samples were collected, homogenized in PBS plus 0.01% NaN_3_ using the Omni Bead Ruptor 24, and centrifuged twice (800 x *g* followed by 10000 x *g*) to remove debris and microbes. Supernatants were filtered through a 0.2 μm filter and Amicon Ultra-0.5 centrifugal filter unit (Millipore Sigma), then analyzed for ATP levels using the ENLITEN ATP Assay System Bioluminescence Detection Kit (Promega) according to the manufacturer’s instructions.

#### Parabiosis

The lateral aspects of CD45.1 mice (Jackson, #002014; left, donor) and CD45.2 *Ccr2^-/-^* mice (Jackson, #004999; right, recipient) were shaved, and matching skin incisions were made from behind the ear to the tail of each mouse, as previously described (Dick *et al*., 2022). The subcutaneous fascia was dissected to create ∼0.5 cm of free skin. The olecranon and knee joints were attached by a mono-nylon 5.0 suture (Ethicon) and the dorsal and ventral skins were attached by continuous suture. Animals recovered with an immediate 0.1mg/kg injection of buprenorphine given subcutaneously. Subcutaneous injections of saline and buprenorphine were given daily for 1 week after the surgery and 3% neomycin antibiotics for 2 weeks. Four-week-old mice were joined for 6 months to 1 year. The CD45.2 recipient mice were analyzed for level of chimerism of CD45.1^+^ cells.

#### Fate-mapping

Tamoxifen was dissolved in corn oil at a concentration of 20 mg/mL by shaking for at least an hour at 55°C and then brought to room temperature. Dissolved tamoxifen was injected intraperitoneally to *Cx3cr1^Cre^*^ERT2^ x *Rosa26*^Td^ mice at 1 mg/kg body weight at day -2 and day 0 prior to the start of the experiment. For *Ccr2^Cre^*^ERT2^ x *Rosa26*^Td^ mice, mice were fed tamoxifen-containing chow (Envigo) for 10 days and then switched back to normal chow during the chase period.

#### Immunofluorescence

Colonic tissues were flushed with 4% formaldehyde, then fixed with 2% formaldehyde 10% sucrose for 1.5 h on ice, followed by an overnight 30% sucrose gradient. Tissues were subsequently embedded in OCT medium (ThermoFisher), flash frozen in 2-methylbutane, and sectioned in 7 μm slices. Sections were blocked and permeabilized for 1 h with blocking/permeabilization buffer (10% BSA, 0.01% Triton X in PBS), washed with PBS, and subsequently stained with antibodies diluted in blocking/permeabilization buffer for 1 h at RT. Sections were mounted with Fluoroshield with DAPI medium (Sigma Aldrich). Slides were imaged at 20X using a Zeiss Axio Imager Z1 and quantified with CellProfiler software (Broad Institute)(McQuin et al., 2018).

#### Adoptive transfer

Leukocytes were isolated from the small intestines of CD45.1/2 *Rorc^+/EGFP^* or CD45.1/2 *Rorc^+/EGFP^* x *Csf2*^-/-^ littermate mice as described above and FACS-sorted for ILC3s using the BD FACSAria II based on Lin^-^ and GFP^+^ expression. Cells were sorted into R-10+ media, checked for purity, then washed with sterile PBS. 10^4^ purified ILC3s were injected intravenously into age- and sex-matched *Rag2*^-/-^*Il2rg*^-/-^ recipient co-housed littermate mice via the retroorbital route. Recipient mice were analyzed 6 weeks later.

### Single-cell RNA sequencing

#### Sample preparation

Colons (n=5 per group) from age- and sex-matched *Cx3cr1^GFP^*^/+^*Ccr2^RFP^*^/+^ WT versus *Csf2*^-/-^ littermate mice were isolated and stripped of epithelium as detailed above, then placed under a Zeiss AxioZoom.V16 fluorescent macroscope for live imaging. Solitary isolated lymphoid tissues (SILTs) were identified based on GFP and RFP expression and isolated using a 1.25 mm biopsy puncher. SILTs and remaining punched out colons were pooled and placed separately into R-10+ media. Samples were digested and enriched for leukocytes as detailed above. Samples were then enriched for CD11b^+^ cells using the EasySep^™^ Mouse CD11b Positive Selection Kit II (StemCell Technologies) as per the manufacturer’s protocol. Purified single cell suspensions (>90% purity) were resuspended in R-10+ media for 10x Genomics single-cell RNA sequencing.

#### Library preparation, sequencing, pre-processing, and quality control

Single cell suspensions were prepared and loaded onto the v3 10x Chromium for the generation of sequencing libraries and processing as described by 10x Genomics. CellRanger (10x Genomics) was used to pre-process sequenced cells, align reads, and generate expression matrices. Seurat (v.4.0) was used for all pre-processing, filtering, and downstream analyses (Hao *et al*., 2021). Low-quality cells expressing fewer than 200 genes were removed. Doublets and dead cells were excluded based on high number of genes (>6000) and high percentage (>9%) of transcripts mapping to mitochondrial genes, respectively. Cells with high percentage (>20%) of transcripts mapping to dissociation-associated genes (DAGs), as previously described, were also removed (O’Flanagan et al., 2019).

#### Normalization, dimensionality reduction, clustering, and cell annotation

To remove technical variation, data was normalized using SCTransform, which utilizes negative binomial regression to normalize the data, find variable features, and scale the data (Hafemeister and Satija, 2019). The variance-stabilizing transformation (vst) method in SCTransform was used to select 3000 highly variable features. Mitochondrial gene percentage and number of counts (nCount_RNA) were regressed out. Dimensionality reduction was performed using principal component analysis (PCA), and an elbow plot was used to determine the number of statistically significant PCs for subsequent clustering. FindNeighbors and FindClusters functions were used to perform graph-based clustering. Non-linear dimensionality reduction and visualization was performed using the Uniform Manifold Approximation and Projection (UMAP) method. Clusters were identified and annotated based on differential gene expression testing using the Wilcoxon Rank Sum Test, with the following parameters in the FindAllMarkers function: min.pct=0.25, logFC threshold=0.25, adjusted p-value<0.05. For heatmaps, each cluster was downsampled to 50 cells for visualization, showing the top 30 differentially expressed genes of each cluster.

Macrophage and monocyte clusters were identified based on expression of MP markers (*C1qa*, *Csf1r*, *Cx3cr1*, and *Adgre1*) and absence of DC markers (*Flt3*, *Dpp4*, *Zbtb46*). These clusters were further subsetted (using the “subset” function), and normalization, dimensionality reduction, and clustering were re-performed as described above to obtain specific MP and monocyte clusters. Clusters were identified, annotated, and visualized as described above.

#### Differential gene expression

To compare gene expression of MPs between wild-type (WT) versus *Csf2*^-/-^ (KO) LP and TLO, the “subset” function was used to separate each cluster from each dataset. The FindMarkers function (min.pct = 0.25, logFC threshold = 0.25, adjusted p value < 0.05) was used to compute differentially upregulated and downregulated genes for each cluster in KO relative to WT of each region (e.g. KO LP relative to WT LP, KO TLO relative to WT TLO). Resulting genes were used for subsequent pathway enrichment analysis, as indicated.

#### Pathway enrichment analysis

gProfiler functional profiling (https://biit.cs.ut.ee/gprofiler/gost) was used to measure over-representation of target gene list against the annotated gene database of Gene Ontology (GO; http://www.geneontology.org). Enriched biological processes of GO (BP, 2019) and enriched KEGG pathways were identified and ordered based on enrichment scores (- log10 of the adjusted p value).

#### Single-cell trajectory analysis

The R package Monocle 3 was used to assess cell trajectories (Cao et al., 2019; Qiu et al., 2017; Trapnell et al., 2014). Data previously analyzed with Seurat (v.4.0), as described above, were imported into Monocle 3 for re-clustering. Briefly, highly variable genes imported from the Seurat analysis were used for PCA dimensionality reduction, followed by UMAP non-linear dimensionality reduction and subsequent clustering using Leidan community detection (https://arxiv.org/abs/1802.03426). The number of Monocle clusters were similar to Seurat clusters. This method also generates ‘partitions’ representing groups corresponding to separate trajectories. Cell trajectory was assessed using the “learn_graph” function, which uses the DDRTree method to learn tree-like trajectories and further reduce dimensionality. Data were visualized with UMAP embeddings and trajectories derived within Monocle and overlaid with Seurat clusters.

#### Quantification and statistical analysis

Statistical analysis of non-sequencing data was performed with the GraphPad Prism software (GraphPad), with statistical tests detailed in the figure legends. All data are shown as mean ± SEM.

## References

Achuthan, A.A., Lee, K.M.C., and Hamilton, J.A. (2021). Targeting GM-CSF in inflammatory and autoimmune disorders. Semin Immunol 54, 101523. 10.1016/j.smim.2021.101523.

Aganna, E., Martinon, F., Hawkins, P.N., Ross, J.B., Swan, D.C., Booth, D.R., Lachmann, H.J., Bybee, A., Gaudet, R., Woo, P., et al. (2002). Association of mutations in the NALP3/CIAS1/PYPAF1 gene with a broad phenotype including recurrent fever, cold sensitivity, sensorineural deafness, and AA amyloidosis. Arthritis Rheum 46, 2445–2452. 10.1002/art.10509.

Amorim, A., De Feo, D., Friebel, E., Ingelfinger, F., Anderfuhren, C.D., Krishnarajah, S., Andreadou, M., Welsh, C.A., Liu, Z., Ginhoux, F., et al. (2022). IFNgamma and GM-CSF control complementary differentiation programs in the monocyte-to-phagocyte transition during neuroinflammation. Nat Immunol 23, 217–228. 10.1038/s41590-021-01117-7.

Atarashi, K., Nishimura, J., Shima, T., Umesaki, Y., Yamamoto, M., Onoue, M., Yagita, H., Ishii, N., Evans, R., Honda, K., and Takeda, K. (2008). ATP drives lamina propria T(H)17 cell differentiation. Nature 455, 808–812. 10.1038/nature07240.

Bain, C.C., Bravo-Blas, A., Scott, C.L., Perdiguero, E.G., Geissmann, F., Henri, S., Malissen, B., Osborne, L.C., Artis, D., and Mowat, A.M. (2014). Constant replenishment from circulating monocytes maintains the macrophage pool in the intestine of adult mice. Nat Immunol 15, 929–937. 10.1038/ni.2967.

Blaut, M., and Clavel, T. (2007). Metabolic diversity of the intestinal microbiota: implications for health and disease. J Nutr 137, 751S–755S. 10.1093/jn/137.3.751S.

Bleriot, C., Chakarov, S., and Ginhoux, F. (2020). Determinants of Resident Tissue Macrophage Identity and Function. Immunity 52, 957–970. 10.1016/j.immuni.2020.05.014.

Bouladoux, N., Harrison, O.J., and Belkaid, Y. (2017). The Mouse Model of Infection with Citrobacter rodentium. Curr Protoc Immunol 119, 19 15 11-19 15 25. 10.1002/cpim.34.

Cao, J., Spielmann, M., Qiu, X., Huang, X., Ibrahim, D.M., Hill, A.J., Zhang, F., Mundlos, S., Christiansen, L., Steemers, F.J., et al. (2019). The single-cell transcriptional landscape of mammalian organogenesis. Nature 566, 496–502. 10.1038/s41586-019-0969-x.

Chang, P.V., Hao, L., Offermanns, S., and Medzhitov, R. (2014). The microbial metabolite butyrate regulates intestinal macrophage function via histone deacetylase inhibition. Proc Natl Acad Sci U S A 111, 2247–2252. 10.1073/pnas.1322269111.

Chiaranunt, P., Burrows, K., Ngai, L., Cao, E.Y., Liang, H., Tai, S.L., Streutker, C.J., Girardin, S.E., and Mortha, A. (2022). NLRP1B and NLRP3 Control the Host Response following Colonization with the Commensal Protist Tritrichomonas musculis. J Immunol. 10.4049/jimmunol.2100802.

Chiaranunt, P., Burrows, K., Ngai, L., and Mortha, A. (2020). Isolation of mononuclear phagocytes from the mouse gut. Methods Enzymol 632, 67–90. 10.1016/bs.mie.2019.10.004.

Chiaranunt, P., Tai, S.L., Ngai, L., and Mortha, A. (2021). Beyond Immunity: Underappreciated Functions of Intestinal Macrophages. Front Immunol 12, 749708. 10.3389/fimmu.2021.749708.

Chuang, L.S., Villaverde, N., Hui, K.Y., Mortha, A., Rahman, A., Levine, A.P., Haritunians, T., Evelyn Ng, S.M., Zhang, W., Hsu, N.Y., et al. (2016). A Frameshift in CSF2RB Predominant Among Ashkenazi Jews Increases Risk for Crohn’s Disease and Reduces Monocyte Signaling via GM-CSF. Gastroenterology 151, 710–723 e712. 10.1053/j.gastro.2016.06.045.

Chudnovskiy, A., Mortha, A., Kana, V., Kennard, A., Ramirez, J.D., Rahman, A., Remark, R., Mogno, I., Ng, R., Gnjatic, S., et al. (2016). Host-Protozoan Interactions Protect from Mucosal Infections through Activation of the Inflammasome. Cell 167, 444–456 e414. 10.1016/j.cell.2016.08.076.

Dai, X.M., Ryan, G.R., Hapel, A.J., Dominguez, M.G., Russell, R.G., Kapp, S., Sylvestre, V., and Stanley, E.R. (2002). Targeted disruption of the mouse colony-stimulating factor 1 receptor gene results in osteopetrosis, mononuclear phagocyte deficiency, increased primitive progenitor cell frequencies, and reproductive defects. Blood 99, 111–120. 10.1182/blood.v99.1.111.

Danne, C., Ryzhakov, G., Martinez-Lopez, M., Ilott, N.E., Franchini, F., Cuskin, F., Lowe, E.C., Bullers, S.J., Arthur, J.S.C., and Powrie, F. (2017). A Large Polysaccharide Produced by Helicobacter hepaticus Induces an Anti-inflammatory Gene Signature in Macrophages. Cell Host Microbe 22, 733–745 e735. 10.1016/j.chom.2017.11.002.

De Schepper, S., Verheijden, S., Aguilera-Lizarraga, J., Viola, M.F., Boesmans, W., Stakenborg, N., Voytyuk, I., Schmidt, I., Boeckx, B., Dierckx de Casterle, I., et al. (2018). Self-Maintaining Gut Macrophages Are Essential for Intestinal Homeostasis. Cell 175, 400–415 e413. 10.1016/j.cell.2018.07.048.

Dick, S.A., Wong, A., Hamidzada, H., Nejat, S., Nechanitzky, R., Vohra, S., Mueller, B., Zaman, R., Kantores, C., Aronoff, L., et al. (2022). Three tissue resident macrophage subsets coexist across organs with conserved origins and life cycles. Sci Immunol 7, eabf7777. 10.1126/sciimmunol.abf7777.

Eberl, G., and Lochner, M. (2009). The development of intestinal lymphoid tissues at the interface of self and microbiota. Mucosal Immunol 2, 478–485. 10.1038/mi.2009.114.

Goudot, C., Coillard, A., Villani, A.C., Gueguen, P., Cros, A., Sarkizova, S., Tang-Huau, T.L., Bohec, M., Baulande, S., Hacohen, N., et al. (2017). Aryl Hydrocarbon Receptor Controls Monocyte Differentiation into Dendritic Cells versus Macrophages. Immunity 47, 582–596 e586. 10.1016/j.immuni.2017.08.016.

Greter, M., Lelios, I., Pelczar, P., Hoeffel, G., Price, J., Leboeuf, M., Kundig, T.M., Frei, K., Ginhoux, F., Merad, M., and Becher, B. (2012). Stroma-derived interleukin-34 controls the development and maintenance of langerhans cells and the maintenance of microglia. Immunity 37, 1050–1060. 10.1016/j.immuni.2012.11.001.

Guendel, F., Kofoed-Branzk, M., Gronke, K., Tizian, C., Witkowski, M., Cheng, H.W., Heinz, G.A., Heinrich, F., Durek, P., Norris, P.S., et al. (2020). Group 3 Innate Lymphoid Cells Program a Distinct Subset of IL-22BP-Producing Dendritic Cells Demarcating Solitary Intestinal Lymphoid Tissues. Immunity 53, 1015–1032 e1018. 10.1016/j.immuni.2020.10.012.

Guilliams, M., De Kleer, I., Henri, S., Post, S., Vanhoutte, L., De Prijck, S., Deswarte, K., Malissen, B., Hammad, H., and Lambrecht, B.N. (2013). Alveolar macrophages develop from fetal monocytes that differentiate into long-lived cells in the first week of life via GM-CSF. J Exp Med 210, 1977–1992. 10.1084/jem.20131199.

Guilliams, M., Thierry, G.R., Bonnardel, J., and Bajenoff, M. (2020). Establishment and Maintenance of the Macrophage Niche. Immunity 52, 434–451. 10.1016/j.immuni.2020.02.015.

Hafemeister, C., and Satija, R. (2019). Normalization and variance stabilization of single-cell RNA-seq data using regularized negative binomial regression. Genome Biol 20, 296. 10.1186/s13059-019-1874-1.

Hamada, H., Hiroi, T., Nishiyama, Y., Takahashi, H., Masunaga, Y., Hachimura, S., Kaminogawa, S., Takahashi-Iwanaga, H., Iwanaga, T., Kiyono, H., et al. (2002). Identification of multiple isolated lymphoid follicles on the antimesenteric wall of the mouse small intestine. J Immunol 168, 57–64. 10.4049/jimmunol.168.1.57.

Han, X., Uchida, K., Jurickova, I., Koch, D., Willson, T., Samson, C., Bonkowski, E., Trauernicht, A., Kim, M.O., Tomer, G., et al. (2009). Granulocyte-macrophage colony-stimulating factor autoantibodies in murine ileitis and progressive ileal Crohn’s disease. Gastroenterology 136, 1261–1271, e1261-1263. 10.1053/j.gastro.2008.12.046.

Hao, Y., Hao, S., Andersen-Nissen, E., Mauck, W.M., 3rd, Zheng, S., Butler, A., Lee, M.J., Wilk, A.J., Darby, C., Zager, M., et al. (2021). Integrated analysis of multimodal single-cell data. Cell 184, 3573–3587 e3529. 10.1016/j.cell.2021.04.048.

Hapfelmeier, S., Lawson, M.A., Slack, E., Kirundi, J.K., Stoel, M., Heikenwalder, M., Cahenzli, J., Velykoredko, Y., Balmer, M.L., Endt, K., et al. (2010). Reversible microbial colonization of germ-free mice reveals the dynamics of IgA immune responses. Science 328, 1705–1709. 10.1126/science.1188454.

Hirata, Y., Egea, L., Dann, S.M., Eckmann, L., and Kagnoff, M.F. (2010). GM-CSF-facilitated dendritic cell recruitment and survival govern the intestinal mucosal response to a mouse enteric bacterial pathogen. Cell Host Microbe 7, 151–163. 10.1016/j.chom.2010.01.006.

Imamura, H., Nhat, K.P., Togawa, H., Saito, K., Iino, R., Kato-Yamada, Y., Nagai, T., and Noji, H. (2009). Visualization of ATP levels inside single living cells with fluorescence resonance energy transfer-based genetically encoded indicators. Proc Natl Acad Sci U S A 106, 15651–15656. 10.1073/pnas.0904764106.

Jones, S.A., Chowdhury, F.Z., Fabich, A.J., Anderson, A., Schreiner, D.M., House, A.L., Autieri, S.M., Leatham, M.P., Lins, J.J., Jorgensen, M., et al. (2007). Respiration of Escherichia coli in the mouse intestine. Infect Immun 75, 4891–4899. 10.1128/IAI.00484-07.

Kang, B., Alvarado, L.J., Kim, T., Lehmann, M.L., Cho, H., He, J., Li, P., Kim, B.H., Larochelle, A., and Kelsall, B.L. (2020). Commensal microbiota drive the functional diversification of colon macrophages. Mucosal Immunol 13, 216–229. 10.1038/s41385-019-0228-3.

Knoop, K.A., Gustafsson, J.K., McDonald, K.G., Kulkarni, D.H., Coughlin, P.E., McCrate, S., Kim, D., Hsieh, C.S., Hogan, S.P., Elson, C.O., et al. (2017). Microbial antigen encounter during a preweaning interval is critical for tolerance to gut bacteria. Sci Immunol 2. 10.1126/sciimmunol.aao1314.

Koscso, B., Kurapati, S., Rodrigues, R.R., Nedjic, J., Gowda, K., Shin, C., Soni, C., Ashraf, A.Z., Purushothaman, I., Palisoc, M., et al. (2020). Gut-resident CX3CR1(hi) macrophages induce tertiary lymphoid structures and IgA response in situ. Sci Immunol 5. 10.1126/sciimmunol.aax0062.

Liu, Z., Gu, Y., Chakarov, S., Bleriot, C., Kwok, I., Chen, X., Shin, A., Huang, W., Dress, R.J., Dutertre, C.A., et al. (2019). Fate Mapping via Ms4a3-Expression History Traces Monocyte-Derived Cells. Cell 178, 1509–1525 e1519. 10.1016/j.cell.2019.08.009.

Macpherson, A.J., and Uhr, T. (2004). Induction of protective IgA by intestinal dendritic cells carrying commensal bacteria. Science 303, 1662–1665. 10.1126/science.1091334.

Matheis, F., Muller, P.A., Graves, C.L., Gabanyi, I., Kerner, Z.J., Costa-Borges, D., Ahrends, T., Rosenstiel, P., and Mucida, D. (2020). Adrenergic Signaling in Muscularis Macrophages Limits Infection-Induced Neuronal Loss. Cell 180, 64–78 e16. 10.1016/j.cell.2019.12.002.

McQuin, C., Goodman, A., Chernyshev, V., Kamentsky, L., Cimini, B.A., Karhohs, K.W., Doan, M., Ding, L., Rafelski, S.M., Thirstrup, D., et al. (2018). CellProfiler 3.0: Next-generation image processing for biology. PLoS Biol 16, e2005970. 10.1371/journal.pbio.2005970.

Mempin, R., Tran, H., Chen, C., Gong, H., Kim Ho, K., and Lu, S. (2013). Release of extracellular ATP by bacteria during growth. BMC Microbiol 13, 301. 10.1186/1471-2180-13-301.

Mortha, A., Chudnovskiy, A., Hashimoto, D., Bogunovic, M., Spencer, S.P., Belkaid, Y., and Merad, M. (2014). Microbiota-dependent crosstalk between macrophages and ILC3 promotes intestinal homeostasis. Science 343, 1249288. 10.1126/science.1249288.

Mortha, A.e.a. (2021). Anti–GM-CSF autoantibodies promote a “pre-diseased” state in Crohn’s Disease. MedRxiv.

Moura Silva, H., Kitoko, J.Z., Queiroz, C.P., Kroehling, L., Matheis, F., Yang, K.L., Reis, B.S., Ren-Fielding, C., Littman, D.R., Bozza, M.T., et al. (2021). c-MAF-dependent perivascular macrophages regulate diet-induced metabolic syndrome. Sci Immunol 6, eabg7506. 10.1126/sciimmunol.abg7506.

Muller, P.A., Koscso, B., Rajani, G.M., Stevanovic, K., Berres, M.L., Hashimoto, D., Mortha, A., Leboeuf, M., Li, X.M., Mucida, D., et al. (2014). Crosstalk between Muscularis Macrophages and Enteric Neurons Regulates Gastrointestinal Motility. Cell 158, 1210. 10.1016/j.cell.2014.08.002.

O’Flanagan, C.H., Campbell, K.R., Zhang, A.W., Kabeer, F., Lim, J.L.P., Biele, J., Eirew, P., Lai, D., McPherson, A., Kong, E., et al. (2019). Dissociation of solid tumor tissues with cold active protease for single-cell RNA-seq minimizes conserved collagenase-associated stress responses. Genome Biol 20, 210. 10.1186/s13059-019-1830-0.

Overacre-Delgoffe, A.E., Bumgarner, H.J., Cillo, A.R., Burr, A.H.P., Tometich, J.T., Bhattacharjee, A., Bruno, T.C., Vignali, D.A.A., and Hand, T.W. (2021). Microbiota-specific T follicular helper cells drive tertiary lymphoid structures and anti-tumor immunity against colorectal cancer. Immunity 54, 2812–2824 e2814. 10.1016/j.immuni.2021.11.003.

Patnode, M.L., Guruge, J.L., Castillo, J.J., Couture, G.A., Lombard, V., Terrapon, N., Henrissat, B., Lebrilla, C.B., and Gordon, J.I. (2021). Strain-level functional variation in the human gut microbiota based on bacterial binding to artificial food particles. Cell Host Microbe 29, 664–673 e665. 10.1016/j.chom.2021.01.007.

Perruzza, L., Gargari, G., Proietti, M., Fosso, B., D’Erchia, A.M., Faliti, C.E., Rezzonico-Jost, T., Scribano, D., Mauri, L., Colombo, D., et al. (2017). T Follicular Helper Cells Promote a Beneficial Gut Ecosystem for Host Metabolic Homeostasis by Sensing Microbiota-Derived Extracellular ATP. Cell Rep 18, 2566–2575. 10.1016/j.celrep.2017.02.061.

Qiu, X., Mao, Q., Tang, Y., Wang, L., Chawla, R., Pliner, H.A., and Trapnell, C. (2017). Reversed graph embedding resolves complex single-cell trajectories. Nat Methods 14, 979–982. 10.1038/nmeth.4402.

Sankowski, R., Bottcher, C., Masuda, T., Geirsdottir, L., Sagar, Sindram, E., Seredenina, T., Muhs, A., Scheiwe, C., Shah, M.J., et al. (2019). Mapping microglia states in the human brain through the integration of high-dimensional techniques. Nat Neurosci 22, 2098–2110. 10.1038/s41593-019-0532-y.

Satoh, J., Kino, Y., Asahina, N., Takitani, M., Miyoshi, J., Ishida, T., and Saito, Y. (2016). TMEM119 marks a subset of microglia in the human brain. Neuropathology 36, 39–49. 10.1111/neup.12235.

Satoh-Takayama, N., Vosshenrich, C.A., Lesjean-Pottier, S., Sawa, S., Lochner, M., Rattis, F., Mention, J.J., Thiam, K., Cerf-Bensussan, N., Mandelboim, O., et al. (2008). Microbial flora drives interleukin 22 production in intestinal NKp46+ cells that provide innate mucosal immune defense. Immunity 29, 958–970. 10.1016/j.immuni.2008.11.001.

Schridde, A., Bain, C.C., Mayer, J.U., Montgomery, J., Pollet, E., Denecke, B., Milling, S.W.F., Jenkins, S.J., Dalod, M., Henri, S., et al. (2017). Tissue-specific differentiation of colonic macrophages requires TGFbeta receptor-mediated signaling. Mucosal Immunol 10, 1387–1399. 10.1038/mi.2016.142.

Schulthess, J., Pandey, S., Capitani, M., Rue-Albrecht, K.C., Arnold, I., Franchini, F., Chomka, A., Ilott, N.E., Johnston, D.G.W., Pires, E., et al. (2019). The Short Chain Fatty Acid Butyrate Imprints an Antimicrobial Program in Macrophages. Immunity 50, 432–445 e437. 10.1016/j.immuni.2018.12.018.

Scott, C.L., Zheng, F., De Baetselier, P., Martens, L., Saeys, Y., De Prijck, S., Lippens, S., Abels, C., Schoonooghe, S., Raes, G., et al. (2016). Bone marrow-derived monocytes give rise to self-renewing and fully differentiated Kupffer cells. Nat Commun 7, 10321. 10.1038/ncomms10321.

Sehgal, A., Donaldson, D.S., Pridans, C., Sauter, K.A., Hume, D.A., and Mabbott, N.A. (2018). The role of CSF1R-dependent macrophages in control of the intestinal stem-cell niche. Nat Commun 9, 1272. 10.1038/s41467-018-03638-6.

Shaw, T.N., Houston, S.A., Wemyss, K., Bridgeman, H.M., Barbera, T.A., Zangerle-Murray, T., Strangward, P., Ridley, A.J.L., Wang, P., Tamoutounour, S., et al. (2018). Tissue-resident macrophages in the intestine are long lived and defined by Tim-4 and CD4 expression. J Exp Med 215, 1507–1518. 10.1084/jem.20180019.

Song, C., Lee, J.S., Gilfillan, S., Robinette, M.L., Newberry, R.D., Stappenbeck, T.S., Mack, M., Cella, M., and Colonna, M. (2015). Unique and redundant functions of NKp46+ ILC3s in models of intestinal inflammation. J Exp Med 212, 1869–1882. 10.1084/jem.20151403.

Theurl, I., Hilgendorf, I., Nairz, M., Tymoszuk, P., Haschka, D., Asshoff, M., He, S., Gerhardt, L.M., Holderried, T.A., Seifert, M., et al. (2016). On-demand erythrocyte disposal and iron recycling requires transient macrophages in the liver. Nat Med 22, 945–951. 10.1038/nm.4146.

Trapnell, C., Cacchiarelli, D., Grimsby, J., Pokharel, P., Li, S., Morse, M., Lennon, N.J., Livak, K.J., Mikkelsen, T.S., and Rinn, J.L. (2014). The dynamics and regulators of cell fate decisions are revealed by pseudotemporal ordering of single cells. Nat Biotechnol 32, 381–386. 10.1038/nbt.2859.

Tsuji, M., Suzuki, K., Kitamura, H., Maruya, M., Kinoshita, K., Ivanov, II, Itoh, K., Littman, D.R., and Fagarasan, S. (2008). Requirement for lymphoid tissue-inducer cells in isolated follicle formation and T cell-independent immunoglobulin A generation in the gut. Immunity 29, 261–271. 10.1016/j.immuni.2008.05.014.

Wan, C.K., Oh, J., Li, P., West, E.E., Wong, E.A., Andraski, A.B., Spolski, R., Yu, Z.X., He, J., Kelsall, B.L., and Leonard, W.J. (2013). The cytokines IL-21 and GM-CSF have opposing regulatory roles in the apoptosis of conventional dendritic cells. Immunity 38, 514–527. 10.1016/j.immuni.2013.02.011.

Witmer-Pack, M.D., Hughes, D.A., Schuler, G., Lawson, L., McWilliam, A., Inaba, K., Steinman, R.M., and Gordon, S. (1993). Identification of macrophages and dendritic cells in the osteopetrotic (op/op) mouse. J Cell Sci 104 ( *Pt 4*), 1021–1029. 10.1242/jcs.104.4.1021.

